# A Compositional Model to Assess Expression Changes from Single-Cell Rna-Seq Data

**DOI:** 10.1101/655795

**Authors:** By Xiuyu Ma, Keegan Korthauer, Christina Kendziorski, Michael A. Newton

## Abstract

On the problem of scoring genes for evidence of changes in the distribution of single-cell expression, we introduce an empirical Bayesian mixture approach and evaluate its operating characteristics in a range of numerical experiments. The proposed approach leverages cell-subtype structure revealed in cluster analysis in order to boost gene-level information on expression changes. Cell clustering informs gene-level analysis through a specially-constructed prior distribution over pairs of multinomial probability vectors; this prior meshes with available model-based tools that score patterns of differential expression over multiple subtypes. We derive an explicit formula for the posterior probability that a gene has the same distribution in two cellular conditions, allowing for a gene-specific mixture over subtypes in each condition. Advantage is gained by the compositional structure of the model, in which a host of gene-specific mixture components are allowed, but also in which the mixing proportions are constrained at the whole cell level. This structure leads to a novel form of information sharing through which the cell-clustering results support gene-level scoring of differential distribution. The result, according to our numerical experiments, is improved sensitivity compared to several standard approaches for detecting distributional expression changes.

## Introduction

The ability to measure genome-wide gene expression at single-cell resolution has accelerated the pace of biological discovery. Overcoming data analysis challenges caused by the scale and unique variation properties of single-cell data will surely fuel further advances in immunology (Papalexi and Satija (2017)), developmental biology (Marioni and Arendt (2017)), cancer (Navin (2015)), and other areas (Nawy (2013)). Computational tools and statistical methodologies created for data of lower-resolution (e.g., bulk RNA-seq) or lower dimension (e.g., flow cytometry) guide our response to the data-science demands of new measurement platforms, but they remain inadequate for efficient knowledge discovery in this rapidly advancing domain (Bacher and Kendziorski (2016)).

An important feature of single-cell studies that could be leveraged better statistically is the fact that cells populate distinct, identifiable subtypes determined by lineage history, epigenetic state, the activity of various transcriptional programs, or other distinguishing factors. Extensive research on clustering cells has produced tools for identifying subtypes, including `SC3` (Kiselev et al. (2017)), `CIDR` (Lin, Troup and Ho (2017)) and `ZIFA` (Pierson and Yau (2015)). We hypothesize that such subtype information may be usefully utilized in other inference procedures in order to improve their operating characteristics.

Assessing the magnitude and statistical significance of changes in gene expression associated with changes in cellular condition has been a central statistical problem in genomics. New tools specific to the single-cell RNA-seq data structure, including `MAST` (Finak et al. (2015)), `scDD` (Korthauer et al. (2016)), and `D3E` (Delmans and Hemberg (2016)), have been deployed to address this problem. These tools respond to scRNA-seq characteristics, such as high prevalence of zero counts and gene-level multimodality, but they do not fully exploit cellular-subtype information. Our proposed method measures changes in a gene’s marginal mixture distribution and acquires sensitivity to a variety of distributional effects by how it integrates gene-level data with estimated cellular subtypes. It is implemented in software in the R package `scDDboost`^1^.

Through the compositional model underlying `scDDboost`, subtypes inferred by clustering inform the analysis of gene-level expression. The proposed methodology merges two lines of computation after cell clustering: one concerns patterns of differential expression among the cellular subtypes, and here we take advantage of the powerful `EBseq` method for detecting patterns in negative-binomially-distributed expression data (Leng et al. (2015)). The second concerns the counts of cells in various subtypes; for this we propose a Double-Dirichlet-Mixture distribution to model the pair of multinomial probability vectors for subtype counts in two experimental conditions. Further elements are developed, on the selection of the number of subtypes and on accounting for uncertainty in the cluster output, in order to provide an end-to-end solution to the differential distribution problem. We note that modularity in the necessary elements provides some methodological advantages. For example, improvements in clustering may be used in place of the default clustering without altering the form of downstream analysis. Also, by avoiding Markov chain Monte Carlo, `scDDboost` computations are relatively inexpensive for a Bayesian procedure.

To set the context by way of example, Figure 1 highlights the ability of `scDDboost` to sense subtype composition changes and thus detect subtle gene expression changes between conditions. The three panels on the left compare expression from 91 human stem cells known to be in the G1 phase of the cell cycle, as well as from 76 such cells known to be in the G2/M phase (Leng et al. (2013)) in three genes (BIRC5, HMMR, and CKAP2), which we happen to know from prior studies have differential activity between G1 and G2/M (Li and Altieri (1999); Sohr and Engeland (2008); Dominguez et al. (2016)). Several standard statistical tools applied to the data behind Figure 1 do not find the observed differences in any of these genes to be statistically significant when controlling the false discovery rate (FDR) at 5%, but `scDDboost` does include these genes on its 5% FDR list. Considering prior studies, these subtle distributional changes are probably not false discoveries. The right panel in Figure 1 shows these three among many other genes also known to be involved in cell-cycle regulation but not identified by standard tools as altered between G1 and G2/M at the 5% FDR level. The color panel provides insight into why `scDDboost` has identified these genes. For this data set, six cellular subtypes were identified in the first step of `scDDboost` (colors red, blue, green, and orange are visible). These subtypes have changed in their proportions between G1 and G2/M; there is a lower proportion of red cells and a greater proportion of orange cells in G2/M, for example. These proportion shifts, which are inferred from genome-wide data, stabilize gene-specific statistics that measure changes between conditions in the mixture distribution of expression, and thereby increase power. We note that `scDDboost` agrees with other statistical tools on very strong differential-distribution signals (not shown), but it has the potential to increase power for subtle signals owing to its unique approach to leveraging cell subtype information.

**Fig 1:**
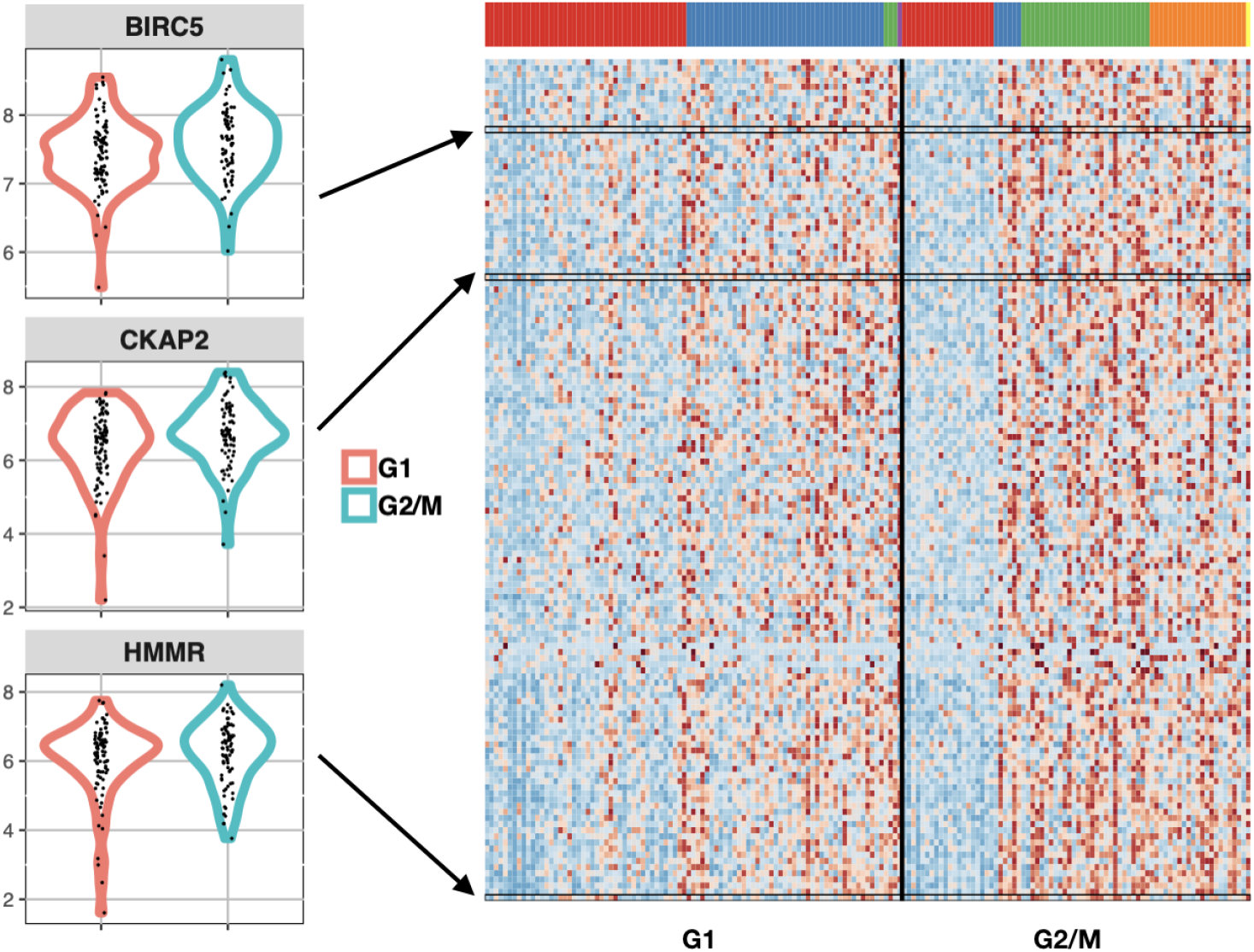
Genes involved in cell-cycle that are identified by scDDboost, but not standard approaches, as differentially distributed between cell-cycle phases G1 and G2/M in human embryonic stem cells. Density estimates on the left show expression data (log2 scale) of three genes identified by scDDboost at 5% FDR, but not similarly identified by MAST, scDD, and DESeq2. Prior studies have shown that the expression of BIRC5, CKAP2, and HMMR is dependent on the phase of cell-cycle, suggesting that these subtle shifts are not false positives. Heatmap (right) shows these three genes among 137 other cell-cycle genes (GO:0007049) identified exclusively by scDDboost, with expression from low (blue) to high (red). Cells (columns) are clustered by their genome-wide expression profiles into distinct cellular subtypes, as indicated by the color panel.

Numerical experiments on both synthetic and published scRNA-seq data bear out the incidental finding in Figure 1, that `scDDboost` has sensitivity for detecting subtle distribution changes. In these experiments we take advantage of `splatter` for generating synthetic data (Zappia, Phipson and Oshlack (2017)) as well as the compendium of scRNA-seq data available through `conquer` (Soneson and Robinson (2017)). Additional numerical experiments show a relationship between `scDDboost` findings and more mechanistic attempts to parameterize transcriptional activation (Delmans and Hemberg (2016)). Finally, we establish first-order asymptotic results for the methodology.

On manuscript organization, we present the modeling and methodology elements in 2, numerical experiments in Section 3, asymptotic analysis in Section 4, and a discussion in Section 5. We relegate some details to an appendix and many others to a Supplementary Material document.

## Results

### Modeling

#### Data structure, sampling model, and parameters

In modeling scRNA-seq data, we imagine that each cell *c* falls into one of *K* > 1 classes, which we think of as subtypes or subpopulations of cells. For notation, *z*_*c*_ = *k* means that cell *c* is of subtype *k*, with the vector *z* = (*z*_*c*_) recording the states of all sampled cells. Knowledge of this class structure prior to measurement is not required, as it will be inferred as necessary from available genomic data. We expect that cells arise from multiple experimental conditions, such as by treatment-control status or some other factors measured at the cell level, but we present our development for the special case of two conditions. Notationally, *y* = (*y*_*c*_) records the experimental condition, say *y*_*c*_ = 1 or *y*_*c*_ = 2. Let’s say condition *j* measures *n*_*j*_ = ∑_*c*_ 1[*y*_*c*_ = *j*] cells, and in total we have *n* = *n*_1_ + *n*_2_ cells in the analysis. The examples in Section 3 involve hundreds to thousands of cells. Further let

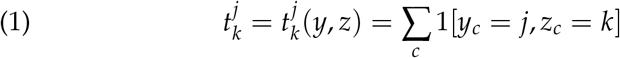

denote the number of cells of subtype *k* in condition *j* and *X*_*g,c*_ denote the normalized expression of gene *g* in cell *c*. This is one entry in a typically large genes-by-cells data matrix *X*. Thus, the data structure entails an expression matrix *X*, a treatment label vector *y*, and a vector *z* of latent subtype labels.

We treat subtype counts in the two conditions, 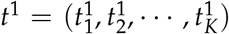 and 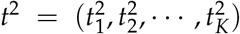, as independent multinomial vectors, reflecting the experimental design. Explicitly,

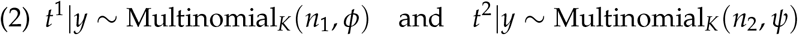

for probability vectors *φ* = (*φ*_1_, *φ*_2_,…, *φ*_*K*_) and *ψ* = (*ψ*_1_, *ψ*_2_,…, *ψ*_*K*_) that characterize the populations of cells from which the *n* observed cells are sampled. This follows from the more basic sampling model: *P*(*z*_*c*_ = *k*|*y*_*c*_ = 1) = *φ*_*K*_ and *P*(*z*_*c*_ = *k*|*y*_*c*_ = 2) = *ψ*_*K*_.

Our working hypothesis, referred to as the *compositional model*, is that any differences in the distribution of expression *X*_*g,c*_ between *y*_*c*_ = 1 and *y*_*c*_ = 2 (i.e., any condition effects) are attributable to differences between the conditions in the underlying composition of cell types; i.e., owing to *φ* ≠ *ψ*. We suppose that cells of any given subtype *k* will present data according to a distribution reflecting technical and biological variation specific to that class of cells, regardless of the condition *y*_*c*_ of the cell. Some care is needed in this, as an overly broad cell subtype (e.g., *epithelial cells*) could have further subtypes that show differential response to some treatment, for example, and so cellular condition (treatment) would then affect the distribution of expression data within the subtype, which is contrary to our working hypothesis. Were that the case, we could have refined the subtype definition to allow a greater number of population classes *K* in order to mitigate the problem of within-subtype heterogeneity. A risk in this approach is that *K* could approach *n*, as if every cell were its own subtype. We find, however, that data sets often encountered do not display this theoretical phenomenon when considering a broad class of within-subtype expression distributions. We revisit the issue in Section 5, but for now, we proceed assuming that cellular condition affects the composition of subtypes but not the distribution of expression within a subtype.

Within the compositional model, let *f*_*g,k*_ denote the sampling distribution of expression measurement *X*_*g,c*_ assuming that cell *c* is from subtype *k*. Then for the two cellular conditions, and at some expression level *x*, the marginal distributions over subtypes are finite mixtures:

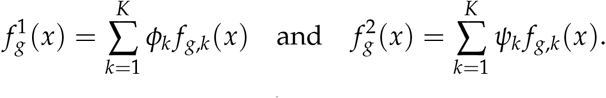

In other words, 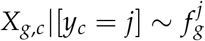 and *X*_*g,c*_|[*z*_*c*_ = *k*, *y*_*c*_ = *j*] *∼ f*_*g,k*_.

We say that gene *g* is *differentially distributed*, denoted DD_*g*_ and indicated by 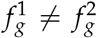, if 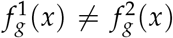 for some *x*, and otherwise it is equivalently distributed (ED_*g*_). Motivated by findings from bulk RNA-seq data analysis, we further set each *f*_*g,k*_ to have a a negative-binomial form, with mean *µ*_*g,k*_ and shape parameter *σ*_*g*_, as in (Leng et al. (2013), Anders and Huber (2010), Love, Huber and Anders (2014) and Chen et al. (2018)). This choice is effective in our numerical experiments though it is not critical to the modeling formulation. The use of mixtures per gene has proven useful in related model-based approaches (e.g., Finak et al. (2015); McDavid et al. (2014); Huang et al. (2018)).

We seek methodology to prioritize genes for evidence of DD_*g*_. Interestingly, even if we have evidence for condition effects on the subtype frequencies, it does not follow that a given gene will have 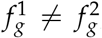; that depends on whether or not the subtypes show the right pattern of *differential expression* at *g*, to use the standard terminology from bulk RNA-seq. For example, if two subtypes have different frequencies between the two conditions (*φ*_1_ ≠ *ψ*_1_ and *φ*_2_ ≠ *ψ*_2_) but the same aggregate frequency (*φ*_1_ + *φ*_2_ = *ψ*_1_ + *ψ*_2_), and also if *µ*_*g,1*_ = *µ*_*g,2*_ then, other things being equal, 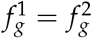 even though *φ* ≠ *ψ*. The fact is so central that we emphasize:

##### Key issue

A gene that does not distinguish two subtypes will also not distinguish the cellular conditions if those subtypes appear in the same aggregate frequency in the two conditions, regardless of changes in the individual subtype frequencies.

We formalize this issue in order that our methodology has the necessary functionality. To do so, first consider the parameter space Θ = {*θ* = (*φ*, *ψ*, *µ*, *σ*)}, where *φ* = (*φ*_1_, *φ*_2_, …, *φ*_*K*_) and *ψ* = (*ψ*_1_, *ψ*_2_, …, *ψ*_*K*_) are as before, where *µ* = {*µ*_*g,k*_} holds all the subtype-and-gene-specific expected values, and where *σ* = {*σ*_*g*_} holds all the gene-specific negative-binomial shape parameters. Critical to our construction are special subsets of Θ corresponding to partitions of the *K* cell subtypes. A single partition, *π*, is a set of mutually exclusive and exhaustive blocks, *b*, where each block is a subset of {1, 2, …, *K*}, and we write *π* = {*b*}. Of course, the set Π containing all partitions *π* of {1, 2, …, *K*} has cardinality that grows rapidly with *K*. We carry along an example involving *K* = 7 cell types, and one three-block partition taken from the set of 877 possible partitions of {1, 2,…, 7} (Figure 2).

**Fig 2:**
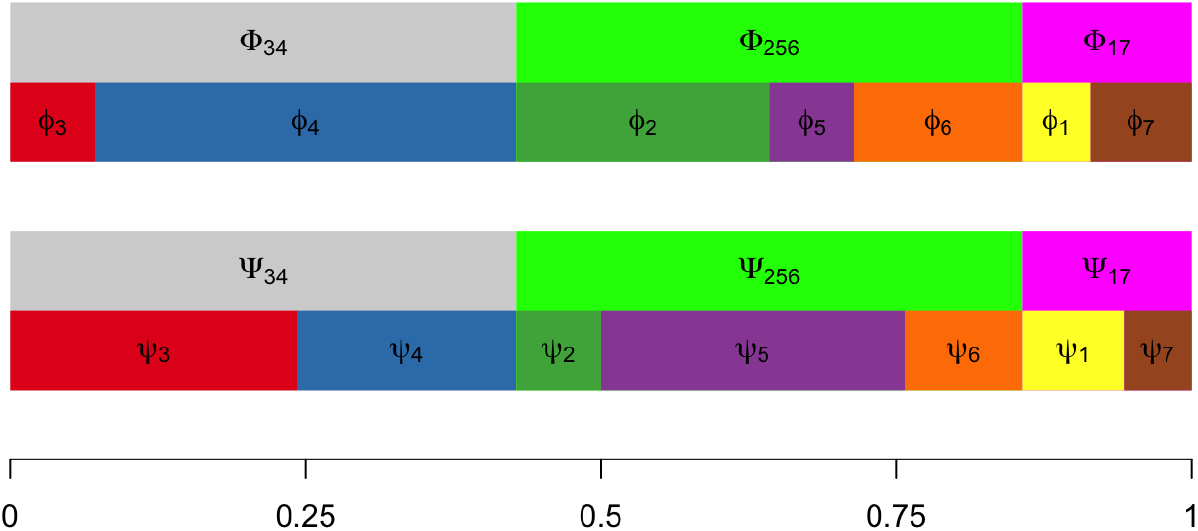
Proportions of *K* = 7 cellular subtypes in two different conditions. Aggregated proportions of subtypes 3 and 4, subtypes 2, 5, and 6, and subtypes 1, and 7 remain same across conditions, while individual subtype frequencies change. Depending on the changes in average expression among subtypes, these frequency changes may or may not induce changes between two conditions in the marginal distribution of some gene’s expression.

For any partition *π* = {*b*}, consider aggregate subtype frequencies

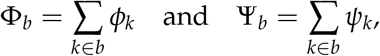

and extend the notation, allowing vectors Φ_*π*_ = {Φ_*b*_: *b* ∈ *π*} and similarly for Ψ_*π*_. Recall the partial ordering of partitions based on refinement, and note that as long as *π* is not the most refined partition (every cell type is in its own block), then the mapping from (*φ*, *ψ*) to (Φ_*π*_, Ψ_*π*_) is many-to-one. Further, define sets

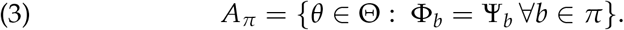

and

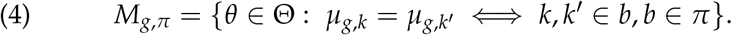

Under *A_π_* there are constraints on cell subtype frequencies; under *M*_*g,π*_ there is equivalence in the gene-level distribution of expression between certain subtypes. These sets are precisely the structures needed to address differential distribution DD_*g*_ (and it complement, equivalent distribution, ED_*g*_) at a given gene *g*, since:

#### Theorem 1.

*Let C*_*g,π*_ = *A*_*π*_ ⋂ *M*_*g,π*_. *For partitions π*_1_ ≠ *π*_2_, 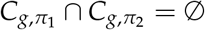. *Further, at any gene g, equivalent distribution is*

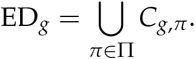

With additional probability structure on the parameter space, we immediately obtain from Theorem 1 a formula for local false discovery rates:

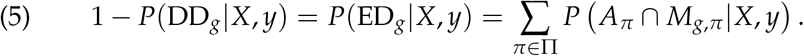

Local false discovery rates are important empirical Bayesian statistics in large-scale testing (Efron (2007); Muralidharan (2010); Newton et al. (2004)). For example, the conditional false discovery rate of a list of genes is the arithmetic mean of the associated local false discovery rates. The partition representation guides the construction of a prior distribution (Section 2.3) and a model-based method (Section 2.2) for scoring differential distribution. Setting the stage, Figure 3 shows the dependency structure of the proposed compositional model and the partition-reliant prior specification.

**Fig 3:**
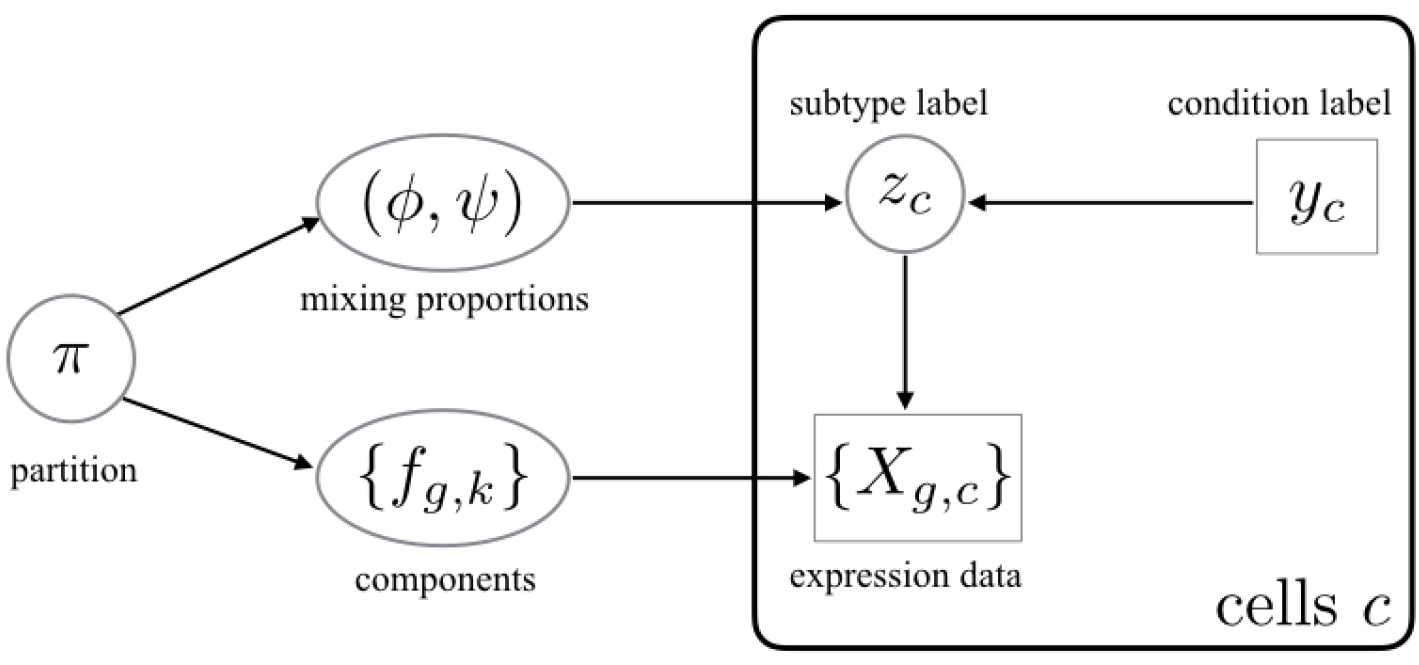
Directed acyclic graph structure of the compositional model and partition-reliant prior. The plate on the right side indicates i.i.d. copies over cells *c*, conditionally on mixing proportions and mixing components. Observed data are indicated in rectangles/squares, and unobserved variables are in circles/ovals.

Key to computing the gene-specific local false discovery rate *P*(ED_*g*_|X, *y*) is evaluating probabilities *P*(*A*_*π*_ ⋂ *M*_*G,π*_|*X,y*). The dependence structure (Figure 3) implies a useful reduction of this quantity, at least conditionally upon subtype labels *z* = (*z*_*c*_). For each subtype partition *π* and gene *g*,

#### Theorem 2.

*P*(*A*_*π*_⋂*M*_*g,π*_|*X,y,z*) = *P*(*A*_*π*_|*y,z*)*P*(*M*_*g,π*_|*X,z*).

In what follows, we develop the modeling and computational elements necessary to efficiently evaluate inference summaries (5) taking advantage of Theorems 1 and 2. Roughly, the methodological idea is that subtype labels *z* have relatively low uncertainty, and may be estimated from genomewide clustering of cells in the absence of condition information *y* (up to an arbitrary label permutation). The modest uncertainty in *z* we handle through a computationally efficient randomized clustering scheme. Theorem 2 indicates that our computational task then separates into two parts given *z*. On one hand, cell subtype frequencies combine with condition labels to give *P*(*A*_*π*_|*y*, *z*). Then gene-level data locally drive the posterior probabilities *P*(*M*_*g,π*_|*X,z*) that measure differential expression between subtypes. Essentially, the model provides a specific form of information sharing between genes that leverages the compositional structure of single-cell data in order to sharpen our assessments of between-condition expression changes.

#### Method structure and clustering

To infer subtypes, we leverage the extensive research on how to cluster cells using scRNA-seq data: for example, SC3 (Kiselev et al. (2017)), CIDR (Lin, Troup and Ho (2017)), and ZIFA (Pierson and Yau (2015)). We propose distance-based clustering on the full set of profiles in a way that is blind to the condition label vector *y*, in order to have as many cells as possible to inform the subtype structure. We investigated several clustering schemes in numerical experiments and allow flexibility in this choice within the scDDboost software. Associating clusters with subtype labels *ẑ*_*c*_ estimates the actual subtypes *z*_*c*_, and prepares us to use Theorems 1 and 2 in order to compute separate posterior probabilities *P*(*A*_*π*_|*y*, *ẑ*) and *P*(*M*_*g,π*_|X, *ẑ*) that are necessary for scoring differential distribution. The first probability concerns patterns of cell counts over subtypes in the two conditions, and has a convenient closed form within the double-Dirichlet model (Section 2.3). The second probability concerns patterns of changes in expected expression levels among sub-types, and this is also conveniently computed for negative-binomial counts using `EBSeq` (Leng et al. (2013)). Algorithm 1 summarizes how these elements combine to get the posterior probability of differential distribution per gene, conditional on an estimate of the subtype labels.

We invoke *K*-medoids (Kaufman and Rousseeuw (1987)) as the default clustering method in scDDboost, and customize the cell-cell distance by integrating two measures. The first assembles gene-level information by cluster-based-similarity partitioning (Strehl and Ghosh (2003)). Separately at each gene, modal clustering (Dahl (2009) and Supplementary Material Section 2.2.2) partitions the cells, and then we define dissimilarity between cells as the Manhattan distance between gene-specific partition labels. A second measure defines dissimilarity by one minus the Pearson correlation between cells, which is computationally inexpensive, less sensitive to outliers than Euclidean distance, and effective at detecting cellular clusters in scRNA-seq (Kim et al. (2018)). The default clustering in scDDboost combines these two measures by weighted average, with 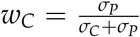 and *w*_*P*_ = 1 − *w*_*C*_, where *w*_*C*_, *σ*_*C*_, *w*_*P*_, *σ*_*P*_ are the weights and standard deviations of cluster-based distance and Pearson-correlation distance, respectively. The software allows other distances; in any case the final distance matrix is denoted *D* = (*d*_*i,j*_).

##### Algorithm 1 scDDBoost-core

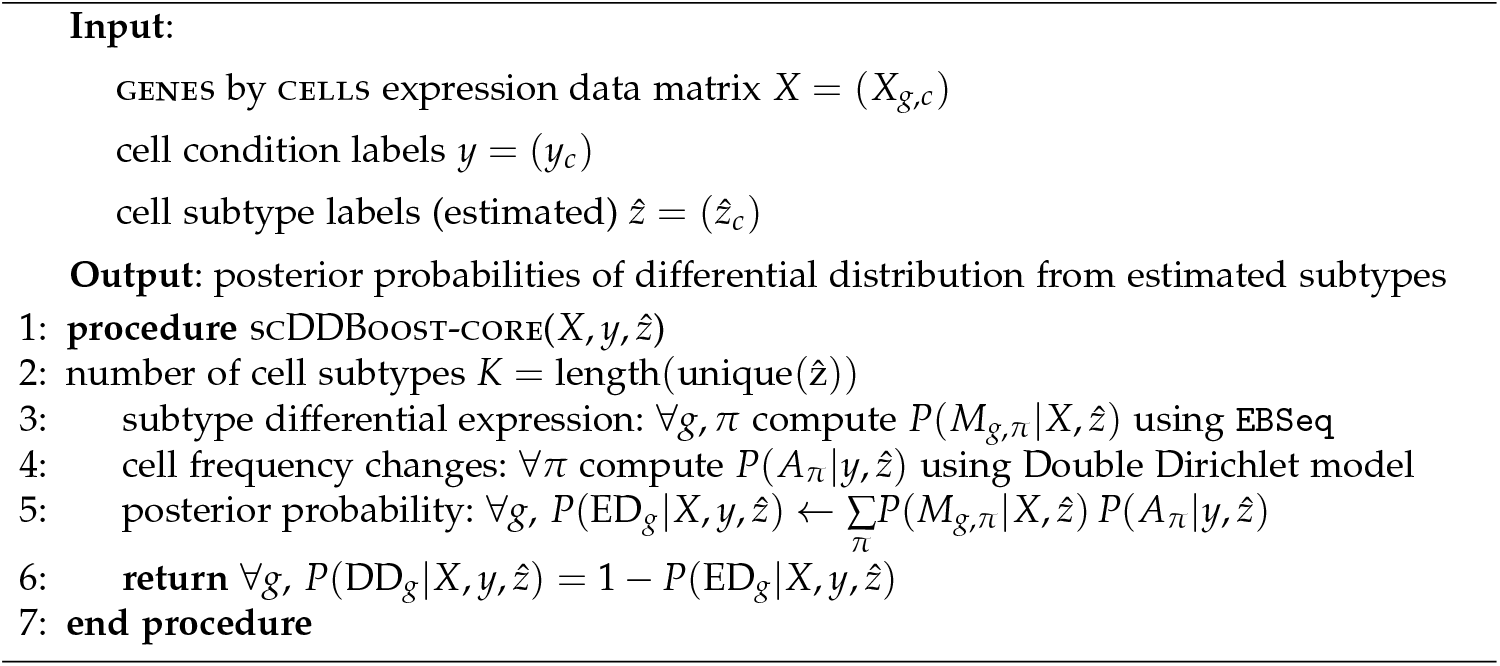

Any clustering method entails classification errors, and so *ẑ*_*c*_ ≠ *z*_*c*_ for some cells. To mitigate the effects of this uncertainty, `scDDboost` averages output probabilities from scDDboost-core over randomized clusterings *ẑ*\*. These are not uniformly random, but rather are generated by applying *K* medoids to a randomized distance matrix *D\** = (*d*_*i,j*_/*w*_*i,j*_), where *w*_*i,j*_ are non-negative weights *w*_*i,j*_ = (*e*_*i*_ + *e*_*j*_), and where (*e*_*i*_) are independent and identically Gamma distributed deviates with shape *â*/2 and rate *â*, and where *â* is estimated from *D*. (Thus *w*_*i,j*_ is Gamma(*â*, *â*) and has unit mean.) The distribution of clusterings induced by this simple computational scheme approximates a Bayesian posterior analysis, as we argue in the Appendix, where we also present pseudo-code for the resulting `scDDboost` Algorithm 2. Averaging over results from randomized clusterings gives additional stability to the posterior probability statistics (Supplementary Figure S10).

Computations become more intensive the larger is the number *K* of cell subtypes. Version 1.0 of `scDDboost` is restricted to *K* ≤ 9; we consider further computational strategies in Section 5. Inferentially, taking *K* to be too large may inflate the false positive rate (Supplementary Figure S11). The approach taken in `scDDboost` is to set *K* using the validity score (Ray and Turi (2000)), which measures changes in within-cluster sum of squares as we increase *K*. Our implementation, in Supplementary Material Section 2.2.4, shows good operating characteristics in simulation.

*P*(*A*_*π*_|*y*, *z*) We introduce the Double Dirichlet Mixture (DDM), which is the partition-reliant prior *p*(*φ*, *ψ*) indicated in Figure 3, in order to derive an explicit formula for *P*(*A*_*π*_|*y*, *z*). We lose no generality here by defining *A*_*π*_ = {(*φ*, *ψ*): Φ_*b*_ = *Ψ*_*b*_ ∀_b_ ∈ π}, rather than as a subset of the full parameter space as in (3). Each *A*_*π*_ is closed and convex subset of the product space holding all possible pairs of length-*K* probability vectors.

We propose a spike-slab-style mixture prior with the following form:

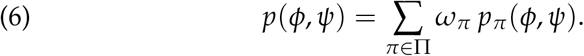

Each mixture component *p*_*π*_(*φ*, *ψ*) has support *A*_*π*_; the mixing proportions *ω*_*π*_ are positive constants summing to one. To specify component *p*_*π*_, notice that on *A*_*π*_ there is a 1-1 correspondence between pairs (*φ*, *ψ*) and parameter states:

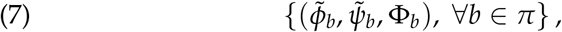

where

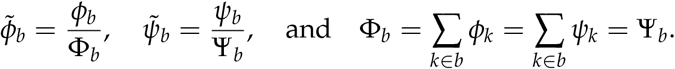

For example, 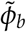 is a vector of conditional probabilities for each subtype given that a cell from the first condition is one of the subtypes in *b*.

We introduce hyperparameters 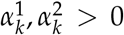 for each subtype *k*, and set 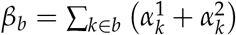 for any possible block *b*. Extending notation, let 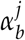 be the vector of 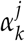 for *k* ∈ *b*, *β*_*π*_ be the vector of *β*_*b*_ for *b* ∈ *π*, *φ*_*b*_ and *ψ*_*b*_ be vectors of *φ*_*K*_ and *ψ*_*K*_, respectively, for *k* ∈ *b*, and Φ_*π*_ and Ψ_*π*_ be the vectors of Φ_*b*_ and Ψ_*b*_ for *b* ∈ *π*. The proposed double-Dirichlet component *p*_*π*_ is determined in the transformed scale by assuming Ψ_*π*_ = Φ_*π*_ and further:

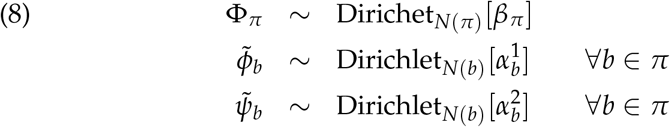

where *N*(*π*) is the number of blocks in *π* and *N*(*b*) is the number of sub-types in *b*, and where all random vectors in (8) are mutually independent. Mixing over *π* as in (6), we write 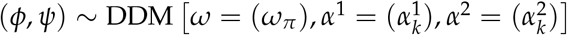.

We record some properties of the component distributions *p*_*π*_:

#### Property 1

In *p*_*π*_(*φ*, *ψ*), *ψ* and *φ* are dependent, unless *π* is the null partition in which all subtypes constitute a single block.

#### Property 2

With *k ∈ b*, marginal means are:

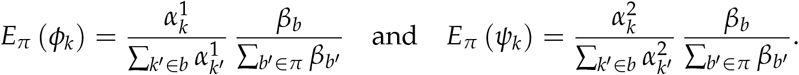

Recall from (1) the vectors *t*^1^ and *t*^2^ holding counts of cells in each sub-type in each condition, computed from *y* and *z*. Relative to a block *b* ∈ *π*, let 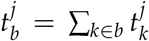, for cell conditions *j* = 1, 2, and, let 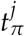 be the vector of these counts over *b* ∈ *π*. The following properties refer to marginal distributions in which (*φ*, *ψ*) have been integrated out of the joint distribution involving (2) and the component *p*_*π*_.

#### Property 3

*t*^1^ and *t*^2^ are conditionally independent given *y*, 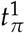 and 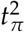.

#### Property 4

For *j* = 1, 2,

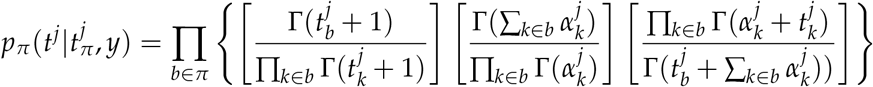

#### Property 5

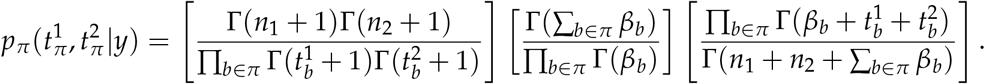

Let’s look at some special cases to dissect this result.

Case 1. If *π* has a single block equal to the entire set of cell types {1, 2,…, *K*}, then 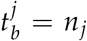 for both *j* = 1, 2, and Property 5 reduces, correctly, to 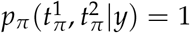. Further,

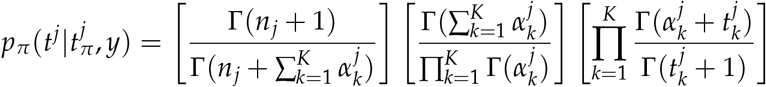

which is the well-known Dirichlet-multinomial predictive distribution for counts *t*^*j*^ (Wagner and Taudes (1986)). E.g, taking 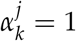 for all types *k* we get the uniform distribution

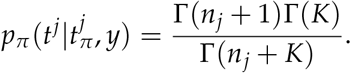

Case 2. At the opposite extreme, *π* has one block *b* for each class *k*, so *φ* = *ψ*. Then 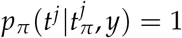, and further, writing *b* = *k*,

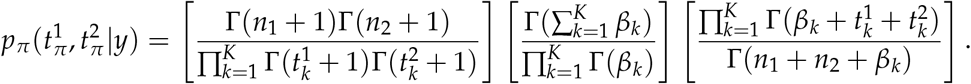

which corresponds to Dirichlet-multinomial predictive distribution for counts *t*^1^ + *t*^2^ since *t*^1^ and *t*^2^ are identical distributed given (*φ*, *ψ*) in this case. These properties are useful in establishing:

#### Theorem 3.

*DDM is conjugate to multinomial sampling of t*^1^ *and t*^2^:

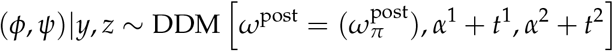

*where*

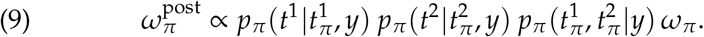

The target probability *P*(*A*_*π*_|*y*, *z*) is an integral of the posterior distribution in Theorem 3. To evaluate it, we need to contend with the fact that sets {*A*_*π*_: *π* ∈ Π} are not disjoint. Relevant overlaps have to do with partition refinement. Recall that a partition *π*^*r*^ is a refinement of a partition *π*^*c*^ if for any *b* ∈ *π*^*c*^ there exists *s* ⊂ *π*^*r*^ such that 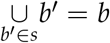. We say *π*^*c*^ coarsens *π*^*r*^ when *π*^*r*^ refines *π*^*c*^. Any partition both refines and coarsens itself, as a trivial case. Generally, refinements increase the number of blocks. If subtype frequency vectors (*φ*, *ψ*) satisfy the constraints in 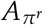 then they also satisfy the constraints of any *π*^*c*^ that coarsens *π*^*r*^: i.e., 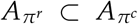. Refinements reduce the dimension of allowable parameter states. For the double-Dirichlet component distributions *p*_*π*_, we find:

#### Property 6

For two partitions 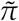 and *π*, 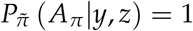 [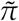 refines *π*].

This supports the main finding of this section:

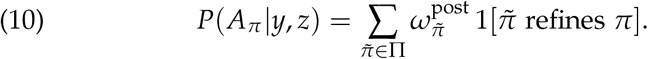

*P*(*M*_*g,π*_|X, *z*) We leverage well-established modeling techniques for transcript analysis, including (Leng et al. (2013), Kendziorski et al. (2003), and Jensen et al. (2009)), which characterize equivalent or differential expression in terms of shared or independently drawn mean effects. Let *X*_*g,b*_ denote the subvector of expression values at gene *g* over cells *c* with *z*_*c*_ = *k* for which subtype *k* is part of block *b* of partition *π*. Conditioning on subtype labels *z* = (*z*_*c*_), we assume that under *M*_*g*,*π*_:

1. *between blocks:* subvectors {*X*_*g,b*_: *b* ∈ *π*} are mutually independent,
2. *within blocks:* for cells mapping to block *b*, observations *X*_*g,c*_ are i.i.d.
3. *mean effects:* for each block *b*, there is a univariate mean, *µ*_*g,b*_, shared by cells mapping to that block. *a priori* these means are i.i.d. between blocks.

These assumptions imply a useful factorization marginally to latent means,

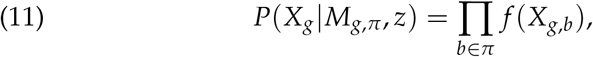

where *f* is a customized density kernel. In our case we use `EBseq` from (Leng et al. (2013)): the sampling distribution of *X*_*g,c*_ is negative binomial, and *f* becomes a particular compound multivariate negative binomial formed from integrating uncertainty in the block-specific means (see Supplementary Material Section 2.2.1). Through its gene-level mixing model, `EBseq` also gives estimates of {*P*(*M*_*g,π*_| z)}: the proportions of genes governed by any of the different patterns *π* of equivalent/differential expression among subtypes. With these estimates and (11) we compute by Bayes’s rule:

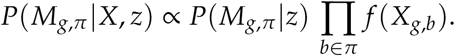

The proportionality is resolved by calculating over all partitions *π*.

### Numerical experiments

#### Synthetic data

We used `splatter` (v. 1.2.0) to generate synthetic scRNA-seq data for which the DD status of genes is known (Zappia, Phipson and Oshlack (2017)), thereby allowing us to measure operating characteristics of scDDboost. Our hypothetical two-condition comparison involved 17421 genes, 10% of which exhibited actual shifts in distribution between two conditions. We entertained 12 different parameter settings encoding these distributional shifts, varying the number of subtypes *K*, the subtype frequency profiles (*φ*, *ψ*), as well as the `splatter`-specific parameters *θ* and *γ* controlling location and scale characteristics of expression levels. These settings cover a range of scenarios we might expect to see in practice. Two replicate data sets were simulated under each parameter setting. Further details are in Supplementary Material Section 3.1.

Figures 4 and 5 summarize the true positive rate and false discovery rate of `scDDboost` compared to three other methodologies: `MAST` (v. 1.4.0), `scDD` (v. 1.2.0), and `DESeq2` (v. 1.18.1). `scDDboost` exhibits very good operating characteristics in this study, as it controls the FDR in all cases while also delivering a relatively high rate of true positives in all cases.

**Fig 4:**
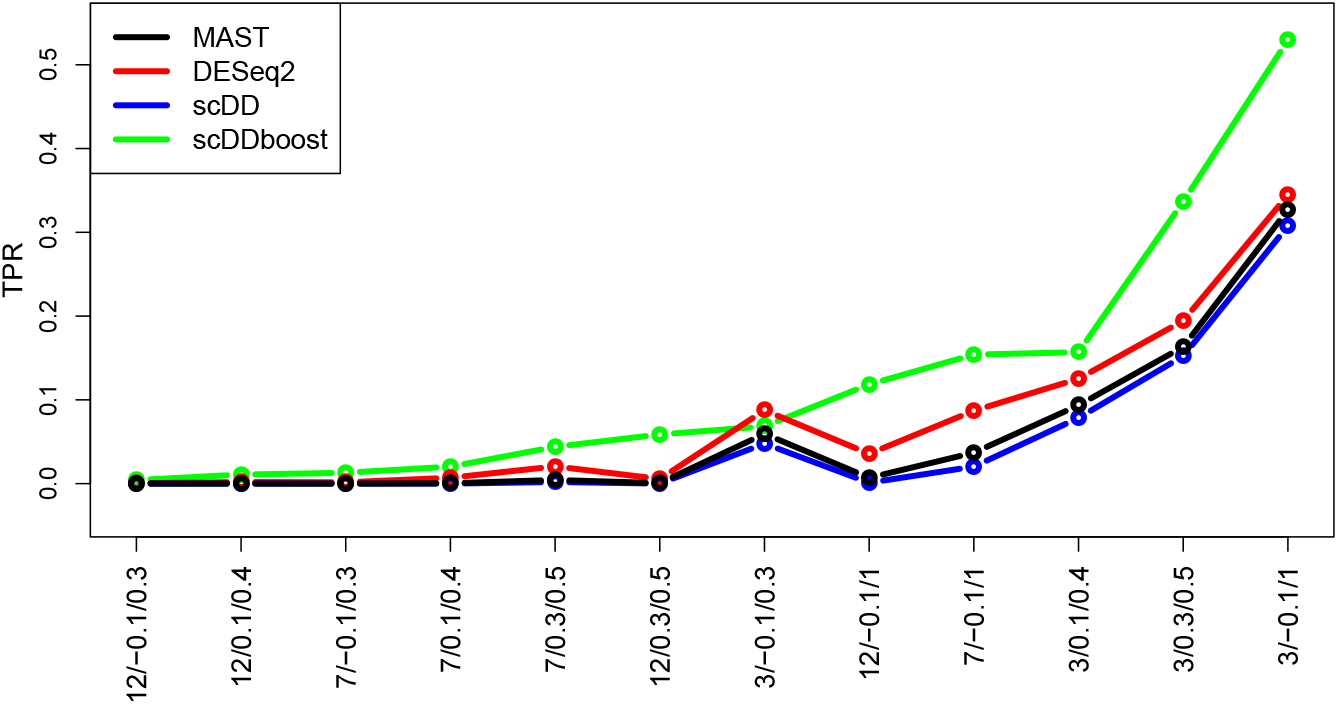
True positive rate (vertical) of four DD detection methods in 12 synthetic-data settings (horizontal). Settings are labeled for *K*/*θ*/*γ* and ranked by scDDboost values. Each method is targeting a 5% false discovery rate (FDR). The plot shows average rates over replicate simulated data in each setting.

**Fig 5:**
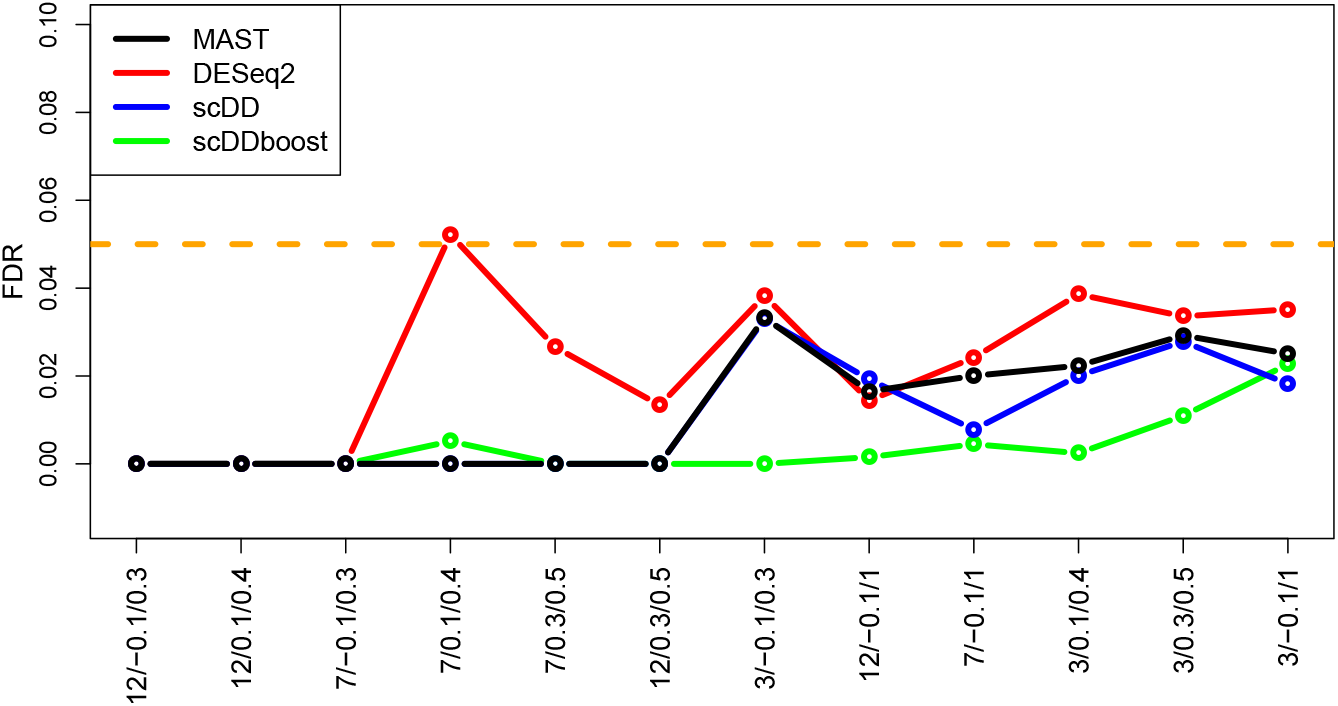
False discovery rate (vertical) of methods in settings (horizontal, same order) from Figure 4

#### Empirical study

We applied `scDDboost` to a collection of previously published data sets that are recorded at `conquer` (Soneson and Robinson (2017)). Though not knowing the truly DD genes, we can examine how `scDDboost` output compares to output from several standard methods. We selected 12 data sets from `conquer` representing different species and experimental settings and involving hundreds to thousands of cells. Appendix Table A1 provides details. Figure 6 compares methods in terms of the size of the reported list of DD genes at the 5% FDR target level. We see a consistently high yield of `scDDboost` among the evaluated methods. For reference, one of these data sets (GSE64016) happens to be the data behind Figure 1, where we know from other information that some of the uniquely identified genes are likely not to be false positives.

**Fig 6:**
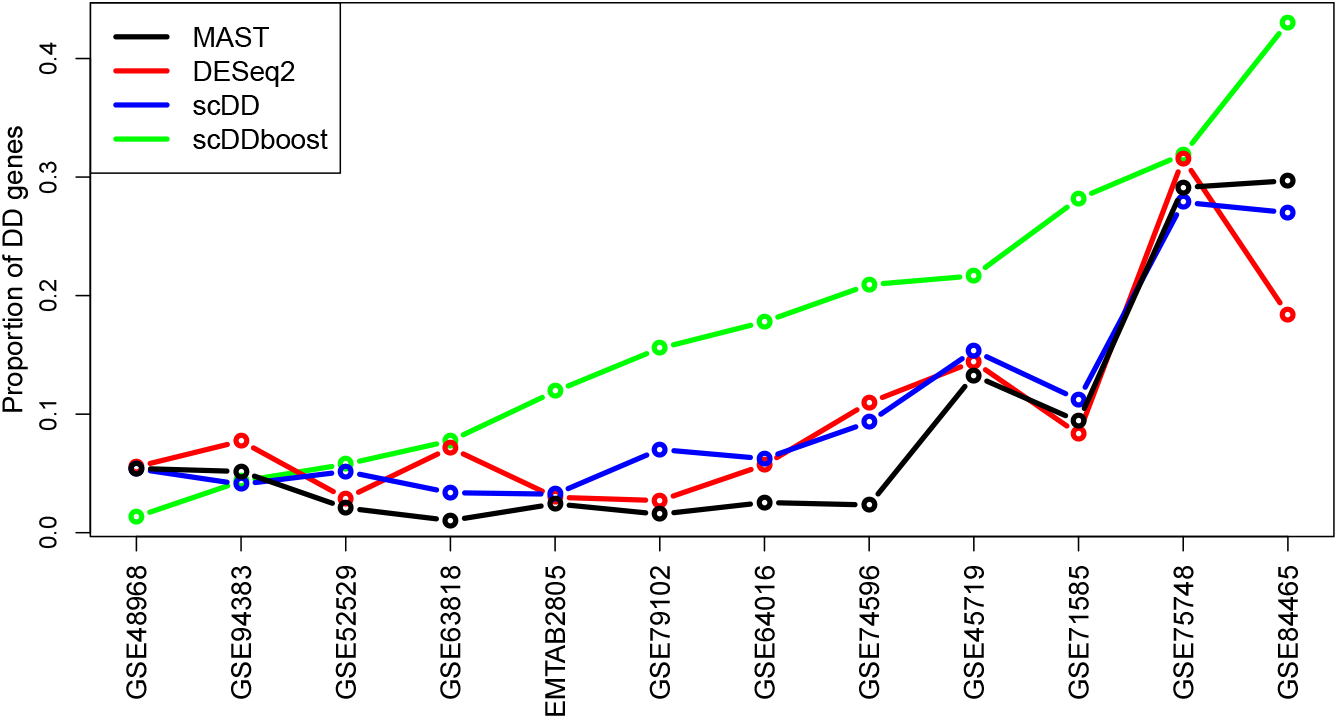
Proportion of DD genes at 5% FDR threshold with respect to total number of genes identified by each method. Ranked by scDDboost list size

To check that the increased discovery rate of `scDDboost` is not associated with an increased rate of false calls, we applied it to a series of random splits of single-condition data sets (Appendix Table A2). Figure 7 confirms a very low call rate in cases where no changes in distribution are expected.

**Fig 7:**
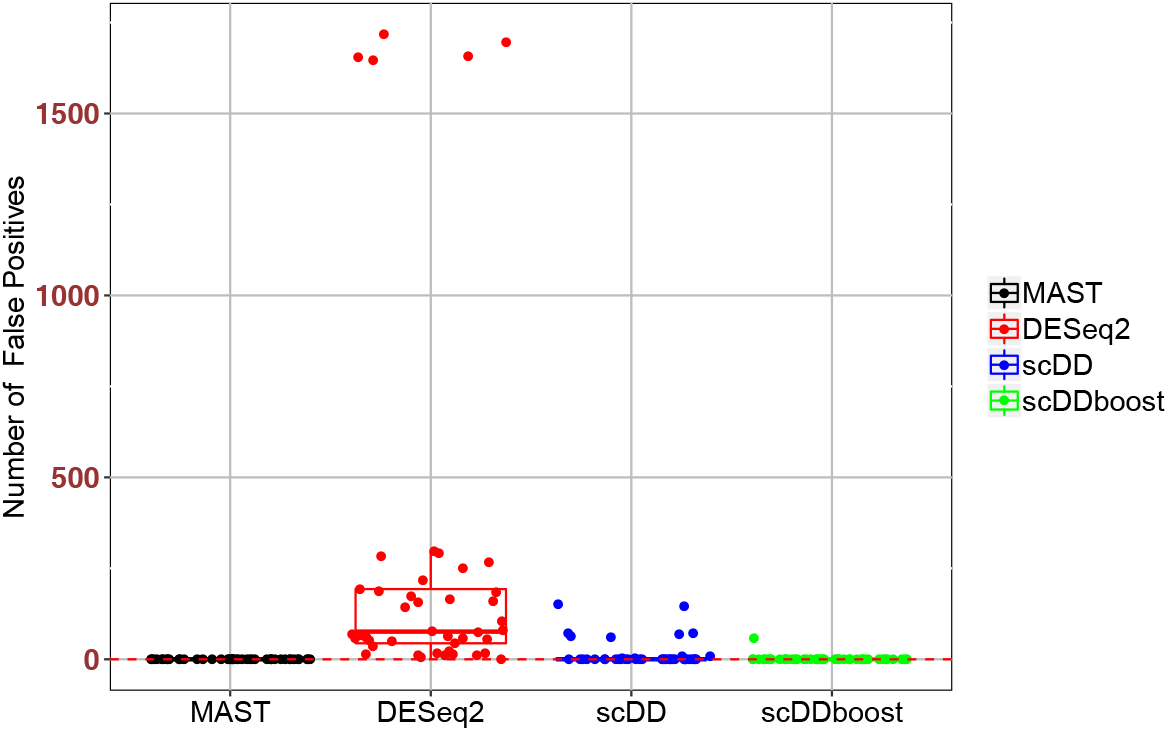
False positive counts at 5% FDR threshold by several methods on 5 random splits of 9 single-condition data sets from Appendix Table A2

We conjecture that `scDDboost` gains power through its novel approach to borrowing strength across genes; i.e., that the genomic data are providing information about cell subtypes and mixing proportions, leaving gene-level data to guide gene-specific mixture components. One way to drill into this idea is to consider how many genes have similar expression characteristics to a given gene. By virtue of the `EBseq` analysis inside `scDDboost`, we may assign each gene to a set of related genes that all have the same highest-probability pattern of equality/inequality of means across the subtypes. Say 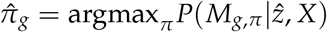. In Figure 8, we show that compared to DD genes commonly identified by multiple methods (blue), the set sizes for genes uniquely identified by `scDDboost` (red) tend to be larger. Essentially, the proposed methodology boosts weak DD evidence when a gene’s pattern of differential expression among cell subtypes matches a large number of other genes.

**Fig 8:**
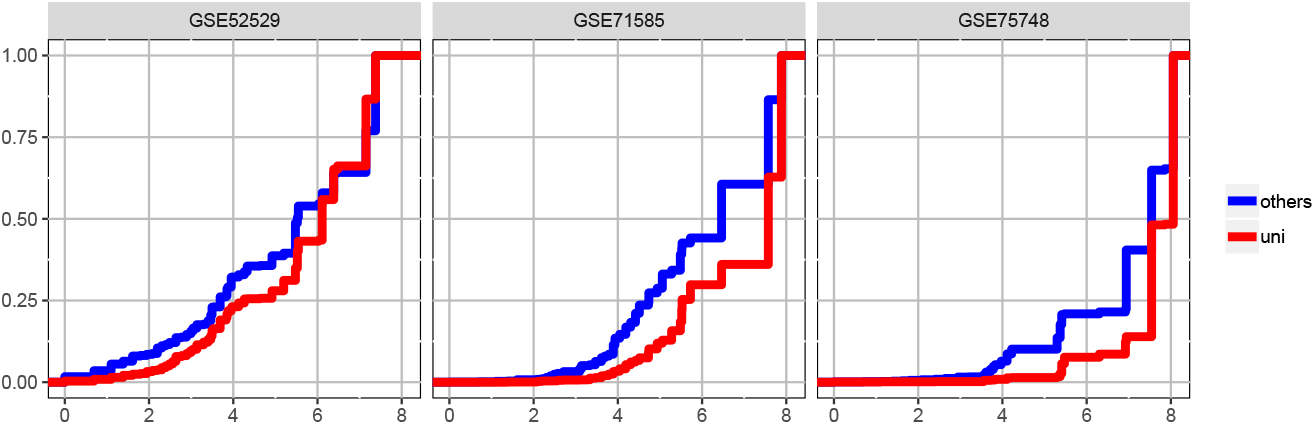
Genes are grouped by their pattern of differential expression across subtypes as inferred by the EBseq computation within scDDboost for three example datasets. Cumulative distribution functions of the log-scale size statistic for all genes identified by scDDboost are plotted; red is the subset uniquely identified by scDDboost; blue are those also identified by the comparison methods (MAST, scDD, or DESeq2). Sets of similarly-patterned genes tend to be larger (horizontal axis, log size) for genes uniquely identified by scDDboost (red) compared to other DD genes (blue), at 5% FDR.

#### Bursting

Transcriptional bursting is a fundamental property of genes, wherein transcription is either negligible or attains a certain probability of activation (Raj and van Oudenaarden (2008)). D3E (Delmans and Hemberg (2016)) is a computationally intensive method for DE gene analysis rooted in modeling the bursting process. It considers transcripts as in the stationary distribution from an experimentally validated stochastic process of single-cell gene expression (Peccoud and Ycart (1995)). Three mechanistic parameters (rate of promoter activation, rate of promoter inactivation, and the conditional rate of transcription given an active promoter) characterize the model, which allow distributional changes between conditions without changing the mean expression level. For genes identified as DD by `scDDboost` in dataset GSE71585, either uniquely or in common with comparison methods, Figure 9 shows changes of these bursting parameters. Interestingly, genes uniquely identified by `scDDboost` are associated with more significant changes between estimated bursting parameters compared to commonly identified genes. This finding and similar findings on other data sets (not shown) provide some evidence that `scDDboost` is able to detect biologically meaningful changes in the expression distribution.

**Fig 9:**
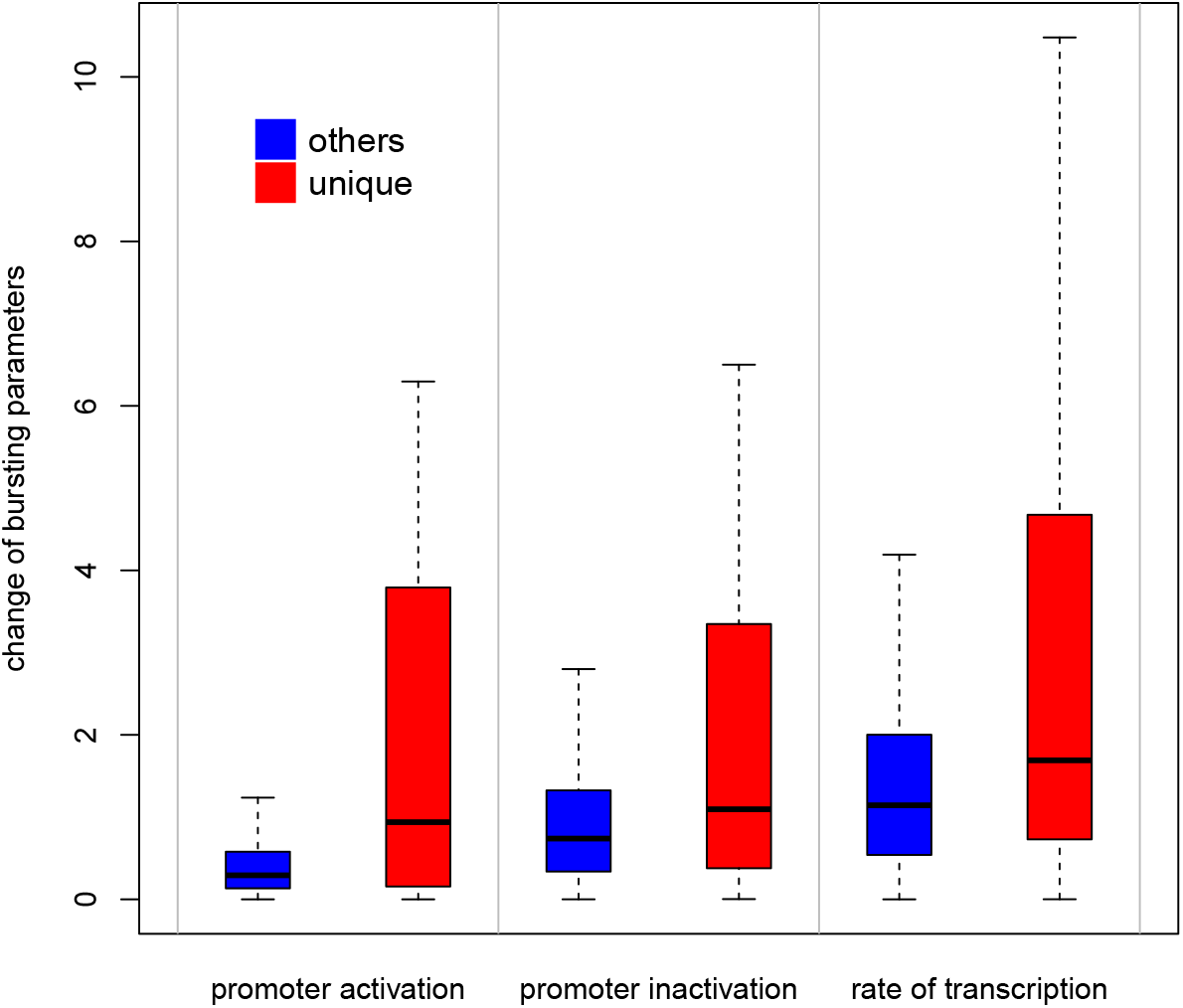
Absolute values of log fold changes of bursting parameters tend to be larger for 1758 genes uniquely identified by scDDboost (red) compare to other 2983 genes (blue) at 5% FDR

#### Time Complexity

Run time complexity of `scDDboost` is dominated by the cost of clustering cells and of running `EBSeq` to measure differences between subtypes. Recall the notation that *n* for number of cells, *G* for number of genes and *K* for number of subtypes. Our distance-based clustering of *n* cells measuring *G* genes requires on the order of *G* × *n*^2^ operations (see Supplementary Material Section 2.2.2). Further, `EBSeq` uses summed counts within each subtype for each gene to compute its density kernel, and there are Bell(*K*) differential patterns to compute, where Bell counts the partitions of *K*. Our implementation `scDDboost` efficiently deals with large *n* under moderate *K*. We have imposed the computational limit *K* ≤ 9 in `scDDboost` (v. 1.0). In a typical case involving 20000 genes and 200 cells, using 50 of randomized distances, `scDDboost` is relatively efficient for *K* ≤ 6 requiring less than 15 CPU minutes on, for example, a quad-core 2.2 GHz Intel Core i7 with 16 Gb of RAM. The same data might require 20 to 40 CPU hours when *K* = 9. In Section 5 we mention some opportunities to improve this speed.

### Asymptotics of the Double Dirichlet Mixture

Summary statistics *P*(*A*_*π*_|*y*, *z*), from Section 2.3, are amenable to a first-order asymptotic analysis that provides further insight into DDM model behavior. The fact that support sets *A*_*π*_ for component distributions *p*_*π*_(*φ*, *ψ*) are not disjoint becomes an important issue. Consider distinct partitions *π*_1_ and *π*_2_ of subtypes {1, 2, …, *K*}, and recall that *N*(*π*) counts the number of blocks in partition *π*. In case *π*_2_ refines *π*_1_, then *N*(*π*_1_) < *N*(*π*_2_), and we also know that 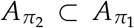, since refinement imposes additional constraints on the pair (*φ*, *ψ*) of probability vectors. If the data-generating state 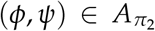, one might ask how posterior probability mass tends to be allocated among the other mixture components whose support sets also contain this state. The question is addressed by the following:

#### Theorem 4.

*Let π*_1_ *and π*_2_ *denote two partitions for which N*(*π*_1_) *< N*(*π*_2_) *and* 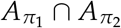 *is non-empty. Let* 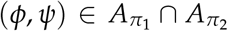 *denote the data generating state for subtype labels z*_1_, *z*_2_, …, *z_n_ given i.i.d. Bernoulli condition labels y*_1_, *y*_2_, …, *y*_*n*_, *and recall the posterior mixing proportions* 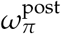 from equation (9) with hyper-parameters 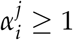 for *i* = 1, …, *K*, *j* = 1, 2. Then

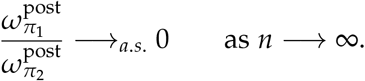

Essentially, mixing mass is transferred to components associated with the most refined partition consistent with a given parameter state. To be precise, let *H*(*φ*, *ψ*) = {*π*: (*φ*, *ψ*) ∈ *A*_*π*_} record all the partitions associated with one state. Typically, there is a most refined partition, *π\** = *π\**(*φ*, *ψ*), such that

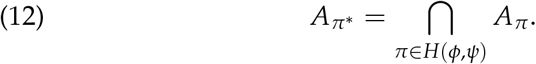

This always happens when *K* ≤ 3. In Supplementary Material Section 4 we characterize the exceptional set of states where (12) does not hold. Notably, if (12) does hold for state (*φ*, *ψ*), then for any *π* ∈ *H*(*φ*, *ψ*), using Theorem 4 and (10), we have

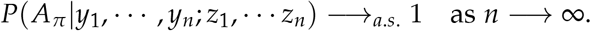

This provides conditions under which we expect good performance for large numbers of cells.

## Discussion

We have presented `scDDboost`, a tool for detecting differentially distributed genes from scRNA-seq data, where transcripts are modeled as a mixture of cellular subtypes. The methodology links established model-based techniques with novel empirical Bayesian modeling and computational elements to provide a powerful detection method showing comparatively good operating characteristics in simulation, empirical, and asymptotic studies.

In the software and numerical experiments we made specific choices, such as to use mixtures of negative binomial components per gene, and to use K-medoids clustering on particular cell-cell distances. These choices have evident advantages, but the model structure and theory developed in Section 2 carry through for other cases. Future experiments could study other formulations within the same schema; for example there may be cell-cell distances that better capture the intrinsic dimensionality of expression programs, including, perhaps distances based on diffusions (Haghverdi, Buettner and Theis (2015)) or the longest-leg path distance (Little, Maggioni and Murphy (2017)). Future experiments could also further assess operating characteristics when the number of cells is very large and the number of reads is relatively small, as may arise with unique molecular identifiers (Chen et al. (2018)). Further, assuming a compositional structure to drive model-based computations may not be restrictive, since it allows great flexibility in the form of each gene/condition-specific expression distribution (as coded, they are finite mixtures of negative binomials).

`EBSeq` currently presents a computational bottleneck for `scDDboost`, since it searches all partitions of *K* and encodes a hyper-parameter estimation algorithm that scales poorly with *K*. Several approximations present themselves that may redress the problem, since, in the mixture model context, only patterns *π* corresponding to relatively probable expression-change patterns over subtypes have a big impact on the final posterior inference. Even resolving this bottleneck there are advantages to having *K* small compared to *n*. Numerical experiments (see Supplement) show increased false discoveries when *K* is over-estimated. But accurate estimation with large *K* would not be expected to provide much improved power, since that depends on accurate estimation of subtypes and their frequencies which relies on *K* being relatively small compared to *n*.

## Appendix

### Proof of Theorem 1

If *θ* ∈ ⋃_*π*∈Π_ [*A*_*π*_ ∩ *M*_*g,π*_], then there exists a partition *π* for which *θ ∈ A*_*π*_ and *θ ∈ M*_*g,π*_. By construction

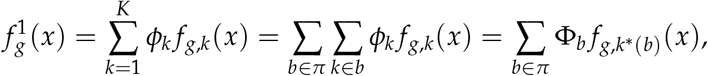

where *k\** (*b*) indexes any component in *b*, since all components in that block have the same component distribution owing to constraint *M*_*g,π*_. Continuing, using the constraint *θ ∈ A*_*π*_,

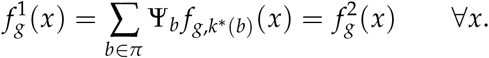

That is, *θ ∈* ED_*g*_.

If *θ ∈* ED_*g*_, then 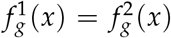 for all *x*. Noting that both are mixtures over the same set of components {*f*_*g,k*_}, let {*h*_*g,l*_: *l* = 1, 2, …, *L*} be the set of distinct components over this set, and so

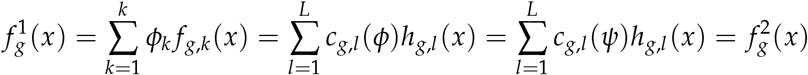

where

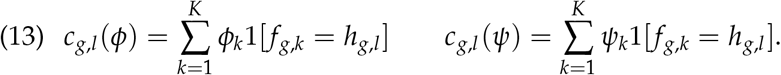

Finite mixtures of distinct negative binomial components are identifiable (Proposition 5 from Yakowitz and Spragins (1968)), and so the equality of 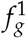 and 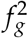 implies *c*_*g,l*_ (*φ*) = *c*_*g,l*_ (*ψ*) for all *l* = 1, 2, …, *L*. Identifying the partition blocks *b*_*l*_ = {*k*: *f*_*g,k*_ = *h*_*g,l*_}, and the partition 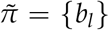, we find 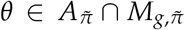. The accumulated probabilities in (13) correspond to 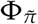 and 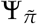, which are equal on 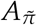.

### Randomizing distances for approximate posterior inference

One way to frame the subtype problem is to suppose that subtype labels *z* = (*z*_*i*_) satisfy *z* = *f* (∆), where ∆ = (*δ*_i,j_) is a *n × n* matrix holding *true*, unobservable distances, such as *δ*_*i,j*_ between cells *i* and *j*, and that *f* is some assignment function, like the one induced by the *K* - medoids algorithm. Then posterior uncertainty in *z* would follow directly from posterior uncertainty in ∆. On one hand, we could proceed via formal Bayesian analysis, say under a simple conjugate prior in which 1/*δ*_*i,j*_ ~ Gamma(*a*_0_, *d*_0_), for hyperparameters *a*_0_ and *d*_0_, and in which the observed distance *d*_*i,j*_ *δ*_*i,j*_ ~ Gamma(*a*_1_, *a*_1_/*δ*_*i,j*_). This would assure that *δ*_*i,j*_ is the expectation of *d*_*i,j*_, with shape parameter *a*_1_ affecting variation of measured distances about their expected values. Not accounting for any constraints imposed by both *D* and ∆ being distance matrices, we would have the posterior distribution 1/*δ*_*i,j*_ D ~ Gamma(*a*_0_ + *a*_1_, *d*_0_ + *a*_1_*d*_*i,j*_). For any threshold *c >* 0, we would find

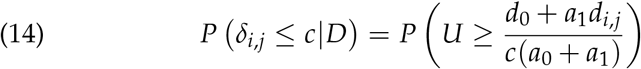

where *U ∼* Gamma(*a*_0_ + *a*_1_, *a*_0_ + *a*_1_)

Alternatively, we could form randomized distances 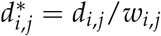 where *w*_*i,j*_ is the analyst-supplied random weight distributed as Gamma(*â*, *â*) as in Section 2.2. Notice that

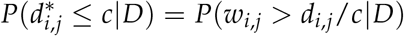

which is also an upper tail probability for a unit-mean Gamma deviate with shape and rate equal to *â*. Comparing to (14), by setting *â* to equal *a*_0_ + *a*_1_, and if *a*_0_ and *d*_0_ are relatively small, we find

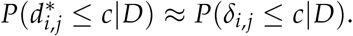

In other words, the randomized distance procedure is providing approximate posterior draws of the underlying distance matrix. In spite of limitations of this procedure for full Bayesian inference, it provides an elementary scheme to account for uncertainty in subtype allocations. Numerical experiments in Supplementary Material make comparisons to a full, Dirichlet-process-based, posterior analysis.

#### Algorithm 2 scDDboost

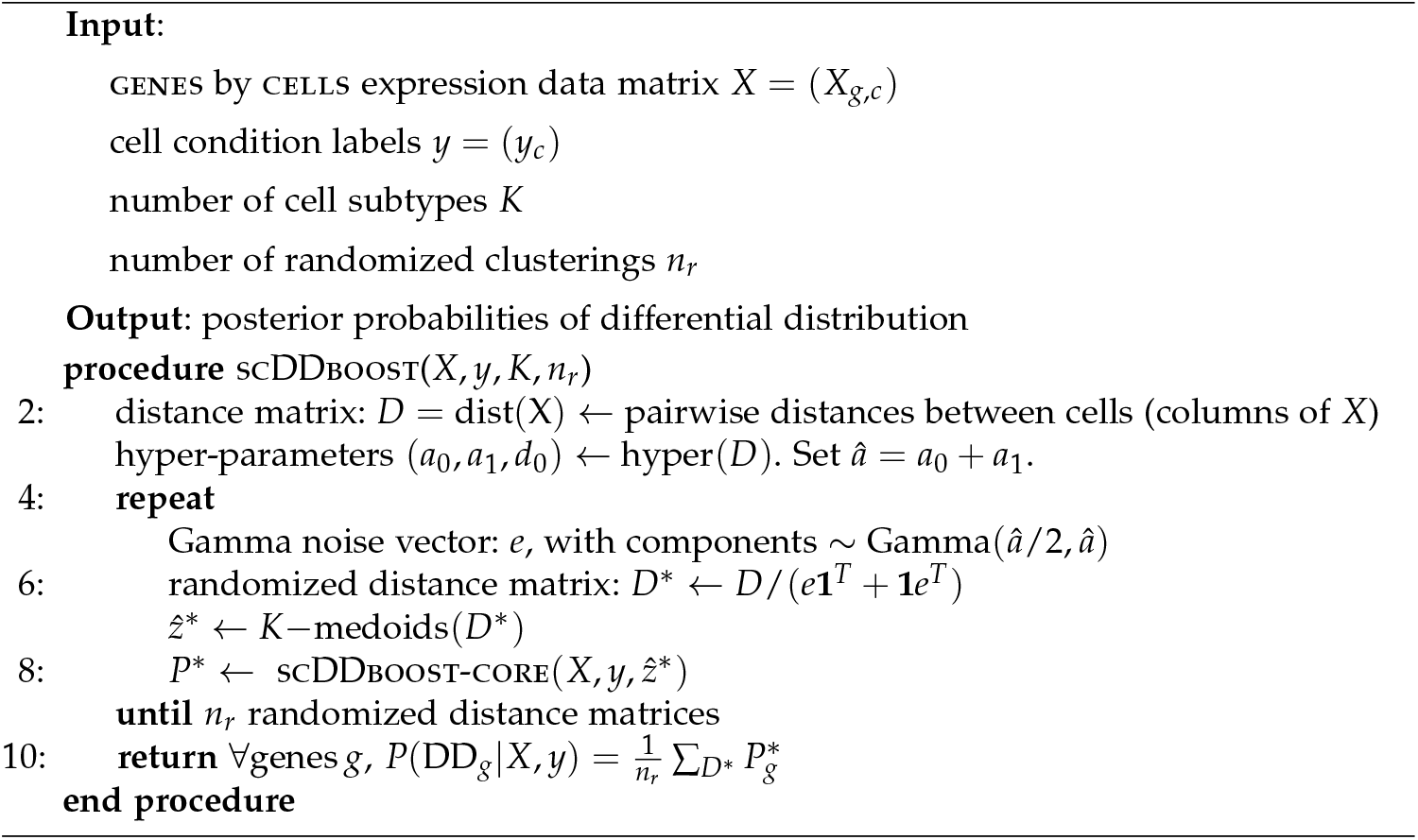

*Pseudo-code*

*Empirical datasets*

**Appendix Table A1.** Data sets used for the empirical study of scDDboost

| Data set | Conditions | # cells | Organism | Ref |
| --- | --- | --- | --- | --- |
| GSE94383 | 0 min unstim vs 75min stim | 186,145 | human | Lane et al. (2017) |
| GSE48968-GPL13112 | BMDC (2h LPS stimulation) vs 6h LPS | 96,96 | mouse | Shalek et al. (2014) |
| GSE52529 | T0 vs T72 | 69,74 | human | Trapnell et al. (2014) |
| GSE74596 | NKT1 vs NTK2 | 46,68 | mouse | Engel et al. (2016) |
| EMTAB2805 | G1 vs G2M | 95,96 | mouse | Buettner et al. (2015) |
| GSE71585-GPL13112 | Gad2tdTpositive vs Cux2tdTnegative | 80,140 | mouse | Tasic et al. (2016) |
| GSE64016 | G1 vs G2 | 91,76 | human | Leng et al. (2015) |
| GSE79102 | patient1 vs patient2 | 51, 89 | human | Kiselev et al. (2017) |
| GSE45719 | 16-cell stage blastomere vs mid blastocyst cell | 50, 60 | mouse | Deng et al. (2014) |
| GSE63818 | Primordial Germ Cells, develop- mental stage: 7 week gestation vs Somatic Cells, developmental stage: 7 week gestation | 40,26 | mouse | Guo et al. (2015) |
| GSE75748 | DEC vs EC | 64, 64 | human | Chu et al. (2016) |
| GSE84465 | neoplastic cells vs non-neoplastic cells | 1000, 1000 | human | Darmanis et al. (2017) |

**Appendix Table A2.** Single-condition data sets used in the random-splitting experiment.

| Data set | Condition | # cells |
| --- | --- | --- |
| GSE63818null | 7 week gestation | 40 |
| GSE75748null | DEC | 64 |
| GSE94383null | T0 | 186 |
| GSE48968-GPL13112null | BMDC (2h LPS stimulation) | 96 |
| GSE74596null | NKT1 | 46 |
| EMTAB2805null | G1 | 96 |
| GSE71585-GPL13112null | Gad2tdTpositive | 80 |
| GSE64016null | G1 | 91 |
| GSE79102null | patient1 | 51 |

## Footnotes

1 http://github.com/wiscstatman/scDDboost/

