## Supplementary material for "A Compositional Model to Assess Expression Changes from Single-Cell Rna-Seq Data"

Version: May 30, 2019

This supplement is organized to match the sectioning of the main document. In summary,

1. Introduction
   - R package
2. Modeling
   - Data Structure, Sampling Model, and Parameters
     Proof of Theorem 2
   - Method Structure and Clustering
     `EBSeq`
     `modalClust`
     Randomized $K-$means
     Selecting $K$
   - Double Dirichlet Mixture
     Proof of Properties 1-6 and Theorem 3
3. Numerical Experiments
   - Synthetic data, `splatter`
   - Empirical study, `conquer` and Null case
   - Robustness
4. Posterior consistency
   - proof of Theorem 4

### 1. Introduction.

1.1. *R package.* We developed a R package `scDDboost` providing clustering of cells, EBSeq and final DD posterior probabilities. Details can be found at github site https://github.com/wiscstatman/scDDboost

### 2. Modeling.

2.1. *Data Structure, Sampling Model, and Parameters.*

Proof of Theorem 2. Recall $\theta = (\phi, \psi, \mu, \sigma)$. Through the graphical structure of our model (Figure 3), given $n_1$ and $n_2$ numbers of cells within each condition, we note that $z^1, z^2$ are multinomial draws, also given $\phi$ and $\psi$. Also given $z$, $X_{g,c}$ is sampled through $\text{NB}(\mu_{g,z_c}, \sigma_g)$, and only depends on $(\mu, \sigma)$. Thus $P(X, y, z|\theta) = P(y, z|\phi, \psi)P(X|z, \mu, \sigma)$, and also $(\mu, \sigma)$ and $(\phi, \psi)$ are independent a priori given $\pi$. By Bayes's rule (and always conditioning on $\pi$),

$$
\begin{aligned}
P(\theta|X, y, z) &\propto P(X, y, z|\theta)P(\theta) \\
P(X, y, z|\theta)P(\theta) &= P(y, z|\phi, \psi)P(X|z, \mu, \sigma)P(\mu, \sigma|z)P(\phi, \psi) \\
P(\phi, \psi|y, z) &\propto P(y, z|\phi, \psi)P(\phi, \psi) \\
P(\mu, \sigma|X, z) &\propto P(X|z, \mu, \sigma)P(\mu, \sigma|z) \\
\text{Thus } P(\theta|X, y, z) &\propto P(\phi, \psi|y, z)P(\mu, \sigma|X, z)
\end{aligned}
$$

It thus follows by integration over the parameter space that $P\left(A_\pi \cap M_{g,\pi}|X, y, z\right) = P\left(A_\pi|y, z\right) P\left(M_{g,\pi}|X, z\right)$. □

2.2. *Method Structure and Clustering.*

2.2.1. *EBSeq.* Here we recall some key elements from Leng et al. (2013) on the model behind `EBSeq`, which we adapt to get $P(M_{g,\pi}|X, z)$. Suppose we have $K$ subtypes, let $X_g^I = X_{g,1}^I, \cdots, X_{g,S_1}^I$ denote transcripts at gene $g$ from subtype $I, I = 1, \cdots, K$. The `EBSeq` model assumes that counts within subtype $I$ are distributed as Negative Binomial: $X_{g,s}^I|r_{g,s}, q_g^I \sim NB(r_{g,s}, q_g^I)$. Due to sample-specific size factor in the raw counts, r is made sample-specific. However, we are dealing with normalized counts rather than raw counts in `EBSeq`, we instead make $r$ shared at gene level across all samples, i.e. $X_{g,s}^I|\sigma_g, q_g^I \sim NB(\sigma_g, q_g^I)$

$$
P(X_{g,s}^I|\sigma_g, q_g^I) = \binom{X_{g,s} + \sigma_g - 1}{X_{g,s}} (1 - q_g^I)^{X_{g,s}^I} (q_g^I)^{\sigma_g}
$$

and $\mu_{g,s}^I = \sigma_g(1 - q_g^I)/q_g^I$; For ease in later deriving the density kernel $f$, we use $q$ rather than $\mu$ to parameterize the NB.

Following Leng et al. (2013), we assume a prior distribution on $q_g^I : q_g^I|\alpha, \beta^g \sim Beta(\alpha, \beta^g)$. The hyperparameter $\alpha$ is shared by the whole genome and $\beta^g$ is gene-specific. We force the size factor to be 1 for all cells and use the same procedure as EBSeq to estimate the shape parameter $\sigma_g$. Namely, we have

1. gene-level sample mean $m_g = \frac{1}{n}\sum_{s=1}^{n} X_{g,s}$, where $n = n_1 + n_2$ is the total number of cells
2. average of sample variances over subtypes $v_g = \frac{1}{K}\sum_{I=1}^{K} v_g^I$.
3. $v_g^I$ is the unadjusted sample variance for subtype $I$, i.e. $v_g^I = \frac{1}{n^I}\sum_{s,z_s=I}(X_{g,s} - m_g^I)^2$ where $m_g^I$ is the sample mean within subtype $I$ and $n^I$ is the number of cells within subtype $I$.

We estimate the pooled over-dispersion rate by $o_g = \frac{v_g}{m_g}$ and obtain $\sigma_g = m_g \frac{o_g}{1-o_g}$ from the first moment of NB. Our aim is to quantify the expression pattern among $K$ groups:

$$M_{g,\pi} = \{\theta \in \Theta : \ \mu_{g,k} = \mu_{g,k'} \iff k, k' \in b, b \in \pi\}.$$

For example, if $K = 3$, there are 5 expression patterns, which may be written equivalently in terms of parameters $q$:

$$\begin{aligned}
&P1 : q_g^1 = q_g^2 = q_g^3 \\
&P2 : q_g^1 = q_g^2 \neq q_g^3 \\
&P3 : q_g^1 \neq q_g^2 = q_g^3 \\
&P4 : q_g^1 = q_g^3 \neq q_g^2 \\
&P5 : q_g^1 \neq q_g^2 \neq q_g^3 \text{ and } q_g^1 \neq q_g^3
\end{aligned}$$

In a pattern where two groups $I$ and $J$ share the same $q_g$ the counts from these groups are essentially pooled: i.e. $X_g^{I,J}|\sigma_g, q_g \sim NB(\sigma_g, q_g)$, $q_g|\alpha, \beta^g \sim \text{Beta}(\alpha, \beta^g)$. The prior predictive function is $f(X_g^{I,J}) = \int_0^1 P(X_g^{I,J}|r_g, q_g) * P(q_g|\alpha, \beta^g) dq_g = \left[\prod_{s=1}^{S} \binom{X_{g,s}+\sigma_g-1}{X_{g,s}}\right] \frac{Beta(\alpha+\Sigma_{s=1}^{S}\sigma_g, \beta^g+\Sigma_{s=1}^{S}X_{g,s})}{Beta(\alpha,\beta^g)}$. Consequently, the prior predictive function for $P1, \cdots, P5$ takes a convenient form if we further treat the distinct $q$'s as independently drawn from the common Beta mixing distribution:

$$\begin{aligned}
h_1^g(X_g) &= f(X_g^{1,2,3}) \\
h_2^g(X_g) &= f(X_g^{1,2}) f(X_g^3) \\
h_3^g(X_g) &= f(X_g^1) f(X_g^{2,3}) \\
h_4^g(X_g) &= f(X_g^{1,3}) f(X_g^2) \\
h_5^g(X_g) &= f(X_g^1) f(X_g^2) f(X_g^3)
\end{aligned}$$

where $h_k^g(X_g) = P(X_g|M_{g,\pi_k}, z)$ for the associated pattern $\pi_k$. Then the marginal distribution of count vector $X_g$ is $\sum_{k=1}^{5} p_k h_k^g(X_g)$, where the mixing mass $p_k = P(M_{g,\pi}|z)$ is shared by all genes. Then, the posterior probability of an expression pattern $k$ is obtained by:

$$\frac{p_k h_k^g(X_g)}{\sum_{l=1}^{5} p_l h_l^g(X_g)}.$$

In the optimization for determining the hyperparameters $(\alpha, \beta^g, p)$, we use EM for the mixing proportions and we use in each cycle a single gradient ascent step for $\alpha$ and $\beta^g$, in contrast to a full root-finding step used by `EBSeq`.

2.2.2. *modalClust.* In this section, we review and extend Dahl's modal clustering procedure (Dahl (2009)). This extension is part of the default cell clustering method of `scDDboost`. It operates on data from one gene at a time, and extends to Poisson-distributed observations the modal-clust procedure.

**Product Partition Model (PPM)**: Let $X = (X_1, X_2, ..., X_n)$ be a vector of data (say at one gene). Given a partition $\pi = \{S_1, \cdots, S_q\}$, where $S_i$ are disjoint subsets of $\{1, 2, \cdots, n\}$ and $\bigcup_{i=1}^{q} S_i = \{1, 2, \cdots, n\}$, a PPM for $X$ entails

$$p(X|\pi) = \prod_{i=1}^{q} f(X_{S_i})$$

where $X_{S_i}$ is the vector of observations corresponding to the items of component $S_i$. The component likelihood $f(X_S)$ is defined for any non-empty component $S$ and can take many forms. The partition $\pi$, which clusters cells, is the parameter we are interested in. Other parameters that may have been involved in the model are integrated out. (Note the partition here has no relation to the partition of subtypes, as, e.g. in Figure 3.)

When the prior distribution for a partition $\pi$ also takes a product form then so does the posterior. We aim to compute the MAP partition (maximizing the posterior $p(\pi|X) \propto p(X|\pi)p(\pi)$) to be used as an initial estimated clustering. Dahl (2009) demonstrated that by some choice of $f$ and prior of $\pi$, we can reduce the time complexity of finding the MAP partition to $O(n^2)$. The crucial condition for $f$ is the **non-overlapping** condition: if $X_{S_1}$ and $X_{S_2}$ are overlapped in the sense that $\min\{X_{S_2}\} < \max\{X_{S_1}\} < \max\{X_{S_2}\}$ or $\min\{X_{S_1}\} < \max\{X_{S_2}\} < \max\{X_{S_1}\}$, let $X_{S_1^*}$ and $X_{S_2^*}$ be the sets of swapping one pair of those overlapped terms and keep the other unchanged. Then $f(X_{S_1})f(X_{S_2}) \leq f(X_{S_1^*})f(X_{S_2^*})$. Here we confirm the non-overlapping condition for Poisson-Gamma observations.

Under the non-overlapping condition of density kernel $f$, the MAP partition $\pi$ satisfies that for any two blocks $b_1, b_2 \in \pi$, either $\max_{i \in b_1}(X_i) \leq \min_{j \in b_2}(X_j)$ or $\min_{i \in b_1}(X_i) \geq \max_{j \in b_2}(X_j)$. Thus we reduce the solution space and reduce the time complexity. In the Poisson-

Gamma model we have:

$$\begin{aligned} X_i|\pi,\lambda &\sim Poisson(X_i|\lambda_1\mathbf{I}\{i\in S_1\}+\cdots+\lambda_q\mathbf{I}\{i\in S_q\}) \\ \pi &\sim p(\pi) \\ \lambda_j &\sim Gamma(\alpha_0,\beta_0) \end{aligned}$$

where $p(\pi) \propto \prod_{i=1}^{q} \eta_0\Gamma(|S_i|)$. Integrate out $\lambda$, $f(X_S)$ is obtained as:

$$f(X_S) = \frac{\beta^\alpha}{(|S|+\beta)^{\sum_{i\in S} X_i+\alpha}} \frac{\Gamma(\sum_{i\in S} X_i+\alpha)}{\Gamma(\alpha)} \frac{1}{\prod_{i\in S} X_i}$$

To apply modal-clustering on Poisson-Gamma model, we need to show the kernel $f(X_S)$ satisfies the non-overlapping condition.

Proof. if $X_{S_1}$ and $X_{S_2}$ are overlapping, without loss of generality, we assume $\min\{X_{S_2}\} < \max\{X_{S_1}\} < \max\{X_{S_2}\}$, and we swap $\max\{X_{S_1}\}$ with $\min\{X_{S_2}\}$ and keep the rest unchanged or we could also swap $\max\{X_{S_1}\}$ with $\max\{X_{S_2}\}$. We denote the new set formed by swap of $\max\{X_{S_1}\}$ with $\min\{X_{S_2}\}$ as $S_1^*$ and $S_2^*$ and swap of $\max\{X_{S_1}\}$ with $\max\{X_{S_2}\}$ as $S_1^{**}, S_2^{**}$ accordingly.

Then we need to show at least one of the following happens

$$f(X_{S_1^*})f(X_{S_2^*}) \geq f(X_{S_1})f(X_{S_2}) \tag{1}$$

$$f(X_{S_1^{**}})f(X_{S_2^{**}}) \geq f(X_{S_1})f(X_{S_2}) \tag{2}$$

Let $a = \max\{X_{S_1}\}$, $b = \min\{X_{S_2}\}$ and $c = \max\{X_{S_2}\}$. $h_1 = \sum_{i\in S_1} X_i - a$ and $h_2 = \sum_{i\in S_2} X_i - b$, $n_1$ and $n_2$ are the number of elements in $S_1$ and $S_2$. Then

$$\begin{gathered} f(X_{S_1^*})f(X_{S_2^*}) \geq f(X_{S_1})f(X_{S_2}) \\ \Longleftrightarrow \\ \frac{\Gamma(h_1+a+\alpha)}{(n_1+\beta)^{h_1+a+\alpha}} \frac{\Gamma(h_2+b+\alpha)}{(n_2+\beta)^{h_2+b+\alpha}} \leq \frac{\Gamma(h_2+a+\alpha)}{(n_2+\beta)^{h_2+a+\alpha}} \frac{\Gamma(h_1+b+\alpha)}{(n_2+\beta)^{h_1+b+\alpha}} \\ \Longleftrightarrow \\ \frac{\Gamma(h_1+a+\alpha)}{\Gamma(h_1+b+\alpha)} \frac{\Gamma(h_2+b+\alpha)}{\Gamma(h_2+a+\alpha)} \leq (\frac{n_1+\beta}{n_2+\beta})^{a-b} \end{gathered}$$

The left hand side of the above formula is $\mathrm{LHS}_1 = \frac{(h_1+b+\alpha)...(h_1+a-1+\alpha)}{(h_2+b+\alpha)...(h_2+a-1+\alpha)}$ by the property of Gamma function and that $X_i$ are integers.

Similarly,

$$f(X_{S_1^{**}})f(X_{S_2^{**}}) \geq f(X_{S_1})f(X_{S_2})$$
$$\Longleftrightarrow$$
$$\frac{\Gamma(h_2+c+\alpha)}{\Gamma(h_2+a+\alpha)}\frac{\Gamma(h_1+a+\alpha)}{\Gamma(h_1+c+\alpha)} \leq (\frac{n_2+\beta}{n_1+\beta})^{c-a}$$

The left hand side of above formula is $\text{LHS}_2 = \frac{(h_2+a+\alpha)...(h_2+c-1+\alpha)}{(h_1+a+\alpha)...(h_1+c-1+\alpha)}$.

If $h_1 \leq h_2$, then $\text{LHS}_1 \leq (\frac{h_1+a-1+\alpha}{h_2+a-1+\alpha})^{a-b}$ and $\text{LHS}_2 \leq (\frac{h_2+c-1+\alpha}{h_1+c-1+\alpha})^{a-b}$.

So if $\frac{h_1+a-1+\alpha}{h_2+a-1+\alpha} \leq \frac{n_1+\beta}{n_2+\beta}$ then (12) holds, if $\frac{h_2+c-1+\alpha}{h_1+c-1+\alpha} \leq \frac{n_1+\beta}{n_2+\beta}$ then (13) holds.

We multiply those two inequalities, and find that $\frac{h_1+a-1+\alpha}{h_2+a-1+\alpha} * \frac{h_2+c-1+\alpha}{h_1+c-1+\alpha} = \frac{h_1+a-1+\alpha}{h_1+c-1+\alpha} * \frac{h_2+c-1+\alpha}{h_2+a-1+\alpha} \leq 1$ as $c > a$ and $h_1 \leq h_2$. But $\frac{n_1+\beta}{n_2+\beta} * \frac{n_1+\beta}{n_2+\beta} = 1$. At least one equality holds, consequently at least one of (12) and (13) holds.

The proof for case $h_1 > h_2$ follows similarly.

□

2.2.3. *Randomized K−means.* In this section, we consider parameters for the distribution of random weights and some properties the induced distribution over cell clusterings. Referring to the appendix in the main paper, to find the value of $a_0, a_1$ and $d_0$, we have the marginal likelihood of $d_{i,j}$

$$P(d_{i,j}|a_0,a_1,d_0) = \frac{\Gamma(a_0+a_1)}{\Gamma(a_0)\Gamma(a_1)}\frac{d_0^{a_0}d_{i,j}^{a_1-1}a_1^{a_1}}{(d_0+a_1*d_{i,j})^{a_0+a_1}}$$

We estimate $d_0$ by treating $d_{i,j} \approx \Delta_{i,j}$ and based on the mean-variance ratio $(\frac{\text{E}(1/\Delta_{i,j})}{\text{Var}(1/\Delta_{i,j})} = d_0)$, $d_0$ can be approximately estimated by moments of $1/d_{i,j}$. Then we obtain $a_0, a_1$ from maximizing marginal density of $d_{i,j}$. The MLE estimators are obtained through `nlminb` function in `R`. One issue that arises is that the default value for tolerance rate of stopping is 1e-10, which yields large value of $a_1 + a_0$ and results in non-randomness of our weighting matrix. To avoid this issue, we set tolerance rate as 1e-3 to obtain moderate deviation from $D$ (Supplementary Figure S1).

We plot the ARI (adjusted RAND index) between randomly generated clusterings to the clustering from the original distances across eight datasets. The boxplots indicate that random weighting is inducing substantial variation in the distribution of cell partitions.

We also check validity of random weighting by comparing it to Dirichlet-process-based clustering (Jara et al. (2011)) on a simulated dataset. We simulate one-dimensional data $X$ from a mixture of 5 normal distributions with different means and same variance $(\mu = (-6,-2,0,2,10), \sigma = 1)$. We compare clustering results between random weighting and Bayesian clustering using the Dirichlet process prior (using `DPpackage`) in terms

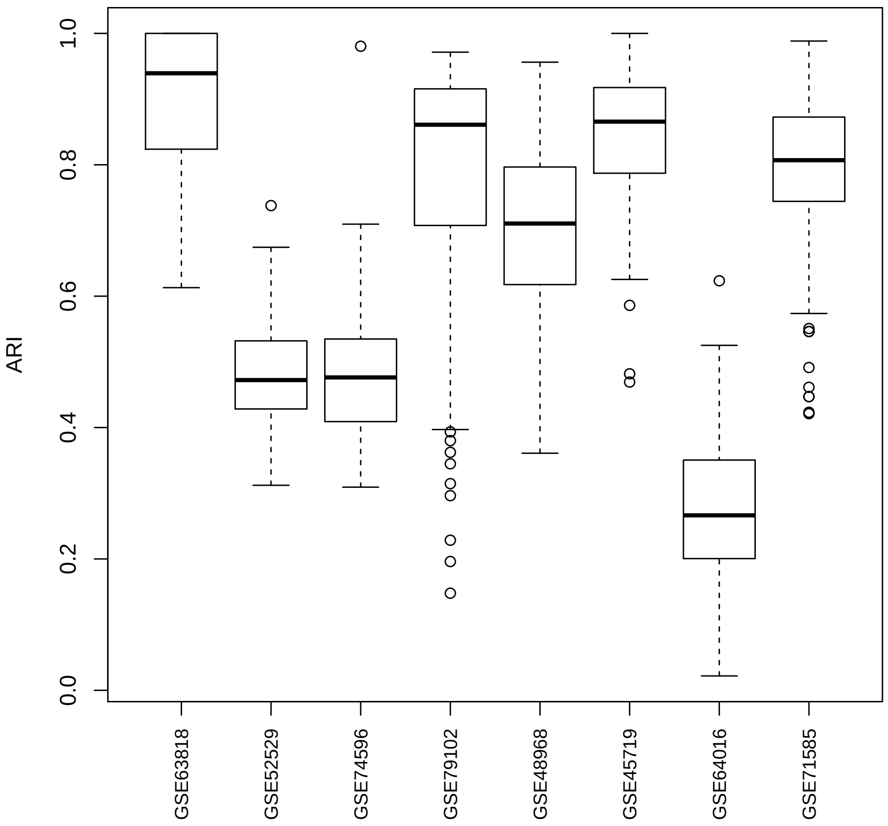


**Supplementary Figure S1:** Adjusted RAND index of clusterings generated by randomizing distances. We investigate the variation of clustering given by random weighting through 8 datasets and each dataset we are using 100 random distances.

of posterior probabilities that two elements belong to the same class given the whole data. We also compare accuracy of the two procedures by looking at the ARI comparing to true class label (Supplementary Figure S2). We find that random weighting scheme closely matches the distributional features of the Dirichlet-process computation, and, in this case, tends to put more mass close to the data-generating partition.

2.2.4. *Selecting K.* In this section, we give the criterion to select the number of subtypes $K$. We implement a procedure inspired by `validity`, as defined in Ray and Turi (2000). We consider a modified `validity`= $\frac{\textbf{intra}}{\textbf{inter}}$, where **intra** $= \frac{1}{N}\sum_{i=1}^{K}\sum_{x\in C_i}||x - z_i||^2$, **inter** $=$ mean$(||z_i - z_j||^2), i, j = 1, 2, ...K$, and $z_i$ is the center (medoid) of cluster $i$. **intra** is the average of distance of a point to the center of its corresponding cluster, which measures the compactness of clusters. **inter** is the average distance between centers, which measures the separation between clusters. In the original paper **inter** was defined as minimum distance between medoids (Ray and Turi, 2000). Here, we instead use the average for smoothness. By minimizing `validity` (contrary to what the name suggests) we aim for

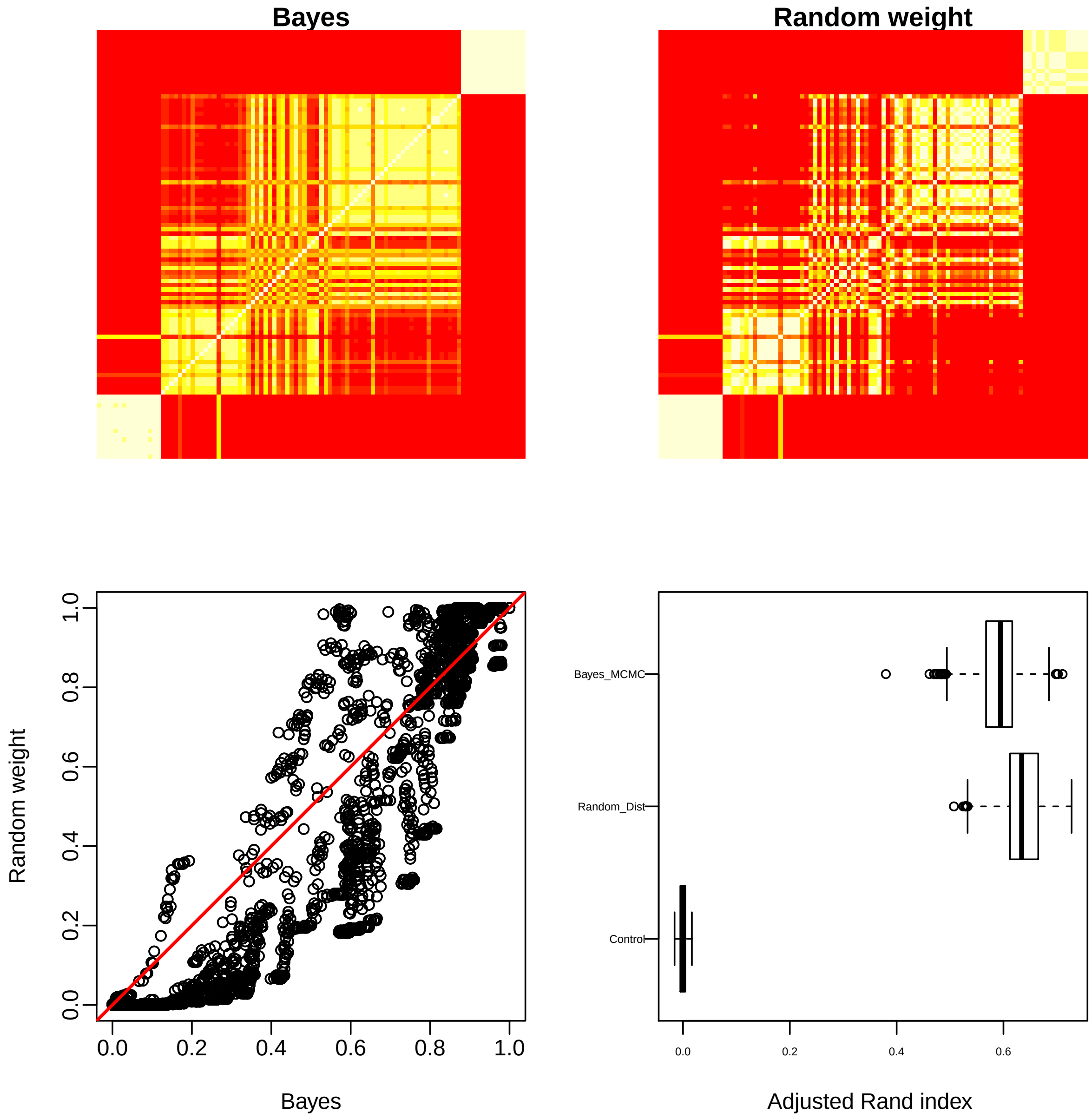


**Supplementary Figure S2:** Comparison between random weighting scheme and Dirichlet-process procedure. Top: heatmap of probabilities that two elements belong to the same class given the whole data. Bottom: scatterplot of these posterior probabilities (left), and adjusted RAND index comparing to the underlying true class label (right).

small intra-cluster distance and large inter-cluster distance. We find empirically that `validity` is monotonic decreasing with $K$ and this trend stabilizes when $K$ is sufficiently big. We select the first $K$ satisfying $\texttt{validity}_K < \epsilon$. We set the default value of $\epsilon$ to be 1, as we found this yields good performance in simulation.

2.3. *Double Dirichlet Mixture.* In this section, we give proofs for the properties of DDM in Section 2.3 of the main paper. Using notation from the main paper, we have density functions:

$$p_\pi(\phi, \psi) = q_\pi(\Phi_\pi, \Psi_\pi) \prod_{b \in \pi} [p(\tilde{\phi}_b)p(\tilde{\psi}_b)]$$

with

$$q_\pi(\Phi_\pi, \Psi_\pi) = \frac{\Gamma(\sum_{b\in\pi} \beta_b)}{\prod_{b\in\pi} \Gamma(\beta_b)} \left[ \prod_{b\in\pi} \Phi_b^{\beta_b - 1} \right] 1\,[\Phi_\pi = \Psi_\pi]$$

and

$$p(\tilde{\phi}_b) = \frac{\Gamma(\sum_{k\in b} \alpha_k)}{\prod_{k\in b} \Gamma(\alpha_k)} \prod_{k\in b} \tilde{\phi}_k^{\alpha_k - 1}, \qquad p(\tilde{\psi}_b) = \frac{\Gamma(\sum_{k\in b} \alpha_k)}{\prod_{k\in b} \Gamma(\alpha_k)} \prod_{k\in b} \tilde{\psi}_k^{\alpha_k - 1}.$$

These serve as key components for proving DDM properties.

Proof of Property 1. When $\phi$ and $\psi$ only satisfy the coarsest constraint: $\sum_{i=1}^K \phi_i = \sum_{i=1}^K \psi_i = 1$, then $\phi$ and $\psi$ are independently Dirichlet distributed. Finer constraints will lead to dependency between $\phi$ and $\psi$ as there is a proper subset $b$ of $\pi$ such that $\sum_{i\in b} \phi = \sum_{i\in b} \psi$, which make $P(\phi|\psi) \neq P(\phi)$. □

Proof of Property 2. By the law of total expectation, $E_\pi(\phi_k) = E_\pi(E_\pi((\phi_k|\Phi_b)) = E_\pi(E_{\tilde{\phi}_b}(\tilde{\phi}_k)) = E_{\tilde{\phi}_b}(\tilde{\phi}_k)E_\Phi(\Phi_b)$ where $b$ is the block containing subtype index $k$. Since $\tilde{\phi}_b \sim \text{Dirichlet}_{N(b)}[\alpha_b^1]$ and $\Phi_\pi \sim \text{Dirichlet}_{N(\pi)}[\beta\pi]$, we have $E_{\tilde{\phi}_b}(\tilde{\phi}_k) = \frac{\alpha_k^1}{\sum_{k'\in b} \alpha_{k'}^1}$, $E_\Phi(\Phi_b) = \frac{\beta_b}{\sum_{b'\in\pi} \beta_{b'}}$ and $E_\pi(\phi_k) = \frac{\alpha_k^1}{\sum_{k'\in b} \alpha_{k'}^1} \frac{\beta_b}{\sum_{b'\in\pi} \beta_{b'}}$ . The case for $E_\pi(\psi_k)$ is similar. □

Proof of Property 3. $t^1/t_\pi^1$ is independent of $t^2/t_\pi^2$ conditioning on $t_\pi^1$ and $t_\pi^2$ by the neutrality property of Dirichlet distribution □

Proof of Property 4. For $j = 1, 2$, let $T_b^j$ be the vector of $t_k^j$ such that $k \in b$. Recall $t_b^j = \sum_{k\in b} t_k^j$. Without loss of generality, we consider the case condition $j = 1$. At the support of $p_\pi$, for different blocks, $T_b^1|\tilde{\phi}_b$ are mutually independent. Then we have factorization:

$$p_\pi(t^1|t_\pi^1, y) = \prod_{b\in\pi} p(T_b^1|t_b^1, y)$$

and right hand side prior predictive function can be obtained by integrating out $\tilde{\phi}_b$. Namely

$$\begin{aligned} p(T_b^1|t_b^1, y) &= \int_{\tilde{\phi}_b} p(T_b^1|\tilde{\phi}_b)p(\tilde{\phi}_b)d\tilde{\phi}_b \\ &= \left\{ \left[ \frac{\Gamma(t_b^j + 1)}{\prod_{k\in b} \Gamma(t_k^j + 1)} \right] \left[ \frac{\Gamma(\sum_{k\in b} \alpha_k^j)}{\prod_{k\in b} \Gamma(\alpha_k^j)} \right] \left[ \frac{\prod_{k\in b} \Gamma(\alpha_k^j + t_k^j)}{\Gamma(t_b^j + \sum_{k\in b} \alpha_k^j))} \right] \right\} \end{aligned}$$

given the prior Dirichlet$[\alpha_b^1]$ of $\tilde{\phi}_b$ and that $p(T_b^1|\tilde{\phi}_b)$ is a multinomial$(\tilde{\phi}_b)$ distribution. □

Proof of Property 5. $t_\pi^1$ and $t_\pi^2$, given the condition label $y$, are independent and identically distributed with $t_\pi^1|\Phi \sim \text{multinomial}(\Phi)$. Thus

$$\begin{aligned} p_\pi(t_\pi^1, t_\pi^2|y) &= \int_\Phi p(t_\pi^1|\Phi)p(t_\pi^2|\Phi)p(\Phi)d\Phi \\ &= \left[\frac{\Gamma(n_1+1)\Gamma(n_2+1)}{\prod_{b\in\pi}\Gamma(t_b^1+1)\Gamma(t_b^2+1)}\right]\left[\frac{\Gamma(\sum_{b\in\pi}\beta_b)}{\prod_{b\in\pi}\Gamma(\beta_b)}\right]\left[\frac{\prod_{b\in\pi}\Gamma(\beta_b+t_b^1+t_b^2)}{\Gamma(n_1+n_2+\sum_{b\in\pi}\beta_b)}\right]. \end{aligned}$$

As prior of $\Phi$ is Dirchlet$[\beta]$ and $n_j = \sum_{b\in\pi} t_b^j$ for $j = 1, 2$. □

To prove Property 6, we need a fact about dimensionality of the intersection of two $A_\pi$'s.

Lemma 1. *If $\pi_2$ is not a refinement of $\pi_1$ then $A_{\pi_1} \cap A_{\pi_2}$ is a lower dimensional subset of $A_{\pi_2}$.*

Proof of Lemma 1. To formalize the problem in linear algebra, we consider the vector space $R^{2K}$, and define a map from block to vector in $R^K$ :$g(b) = v_b$, where $i$th component of $v_b$ is 1 if $i \in b$ and 0 otherwise.

Let $V_1, V_2$ denote the orthogonal space of $\phi - \psi$ when $(\phi, \psi) \in \cap A_{\pi_2}, A_{\pi_2}, A_{\pi_1},$. Notice that $\dim(A_{\pi_1} \cap A_{\pi_2}) = \dim(\phi - \psi) + \dim(\psi) = K - \dim(V_1) + K - 1 = 2K - \dim(V_1) - 1$, $\dim(A_{\pi_2}) = 2K - \dim(V_2) - 1$, $\dim(V_2) = N(\pi_2)$. Assuming $\pi_1 = \{b_1^1, \cdots, b_s^1\}$, and $\pi_2 = \{b_1^2, \cdots, b_t^2\}$. The corresponding vectors are $v_1^1, \cdots, v_s^1$ and $v_1^2, \cdots, v_t^2$. We claim there must be a $b_i^1 \in \pi_1$ whose corresponding vector $v_i^1$ is linear independent with $v_1^2, \cdots, v_t^2$. If not, for every $v_i^1$ there exists $\alpha_1^i, \cdots, \alpha_t^i$ such that

$$v_i^1 = \sum_{j=1}^t \alpha_j^i v_j^2 \qquad (*)$$

If $b_j^2 \cap b_i^1 \neq \emptyset$, then $v_i^1 * v_j^2 > 0$ and we multiply $v_j^2$ on both sides of (*), we obtain $v_i^1 * v_j^2 = \alpha_j^i (v_j^2)^2$, as $v_p^2 * v_q^2 = 0$ if $p \neq q$. This implies $\alpha_j^i > 0$. Consider $x = g(b_j^2 \setminus b_i^1)$. We have $x * v_i^1 = 0$ and multiply $x$ on both sides of (*) to obtain $\alpha_j^i v_j^2 * x = 0$. Thus x must be the zero vector and $b_j^2 \setminus b_i^1 = \emptyset$, which implies $b_j^2 \subset b_i^1$. That is to say when $b_j^2 \cap b_i^1 \neq \emptyset$, $b_j^2$ must be a subset of $b_i^1$. So $b_i^1$ is the union of some blocks in $\pi_2$. This implies $\pi_2$ is a refinement of $\pi_1$, which is a contradiction. Consequently, there exists $b \in \pi_1$ whose $v_b$ is linear independent with $v_{b'}, \forall b' \in \pi_2$. Thus the $\dim(V_1)$ is is at least $N(\pi_2) + 1, \dim(A_{\pi_1} \cap A_{\pi_2}) < \dim(A_{\pi_2})$. □

Proof of Property 6. For any $\pi$, $P(A_\pi, |y, z) = \sum_{\tilde\pi\in\Pi} \int_{A_\pi} \omega_{\tilde\pi}^{\text{post}} d\phi d\psi$, notice the support of $\omega_{\tilde\pi}^{\text{post}}$ is $A_{\tilde\pi}$. By Lemma 1, we know if $\tilde\pi$ does not refine $\pi$, then $\int_{A_\pi} \omega_{\tilde\pi}^{\text{post}} d\phi d\psi$ is an integral on lower dimension set and vanishes. if $\tilde\pi$ refines $\pi$, then $\int_{A_\pi} \omega_{\tilde\pi}^{\text{post}} d\phi d\psi = \int_{A_{\tilde\pi}} \omega_{\tilde\pi}^{\text{post}} d\phi d\psi = \omega_{\tilde\pi}^{\text{post}}$. We have $P(A_\pi, |y, z) = \sum_{\tilde\pi\in\Pi} \omega_{\tilde\pi}^{\text{post}} 1[\tilde\pi \text{ refines } \pi]$. □

PROOF OF THEOREM 3. Recall the DDM prior: $p(\phi,\psi) = \sum_{\pi\in\Pi} p_\pi(\phi,\psi)$. By Bayes's rule $p(\phi,\psi|y,z) \propto p(\phi,\psi,y,z) = \sum_{\pi\in\Pi} p(y,z|\phi,\psi)p_\pi(\phi,\psi)\omega_\pi$ and the 1-1 map from $(\phi,\psi)$ to $(\tilde\phi,\tilde\psi,\Phi)$, we have

$$p(y,z|\phi,\psi)p_\pi(\phi,\psi) = p(y,z|\tilde\phi,\tilde\psi,\Phi_\pi)p(\tilde\phi)p(\tilde\psi)p(\Phi_\pi)$$

when $(\phi,\psi)\in A_\pi$. Let us denote right hand side of the above equation as $U_\pi$, then

$$U_\pi = \omega_\pi A_1 A_2 A_3 \prod_{k=1}^{K} (\tilde\phi_k)^{t_k^1+\alpha_k^1}(\tilde\psi_k)^{t_k^2+\alpha_k^2} \prod_{b\in\pi}(\Phi_b)^{t_b^1+t_b^2+\beta_b},$$

where $A_1$ is the product of normalizing terms from multinomial distribution of $z^1$ and $z^2$, $A_1 = \frac{\Gamma(n_1+1)\Gamma(n_2+1)}{\prod_{j=1}^2\prod_{k=1}^K \Gamma(t_k^j+1)}$, and $A_2$ is the product of normalizing terms from Dirichlet distribution of $\tilde\phi$ and $\tilde\psi$, $A_2 = \frac{\Gamma(\sum_{k=1}^K \alpha_k^1+1)\Gamma(\sum_{k=1}^K \alpha_k^2+1)}{\prod_{j=1}^2\prod_{k=1}^2 \Gamma(\alpha_k^j+1)}$, and $A_3$ is the normalizing term from Dirichlet distribution of $\Phi_\pi$, $A_3 = \frac{\Gamma(\sum_{b\in\pi}\beta_b+1)}{\prod_{b\in\pi}\Gamma(\beta_b+1)}$. Looking at the indices of $\tilde\phi,\tilde\psi$ and $\Phi$, we can decompose $U_\pi$ as a product of three Dirichlet densities with a normalizing term. Namely $U_\pi = C_\pi * f_1 f_2 f_3$, where $f_1 \sim \text{Dirichlet}[\alpha^1+t^1]$, $f_2 \sim \text{Dirichlet}[\alpha^2+t^2]$ and $f_3 \sim \text{Dirichlet}[\beta+t^1+t^2]$. Considering the normalizing factors for densities $f_1, f_2$ and $f_3$, and multiplying them with $A_1, A_2$ and $A_3$, we have $C_\pi = p_\pi(t^1|t_\pi^1,y)\, p_\pi(t^2|t_\pi^2,y)\, p_\pi(t_\pi^1,t_\pi^2|y)\omega_\pi$. Consequently, we have

$$(\phi,\psi)|y,z \sim \text{DDM}\left[\omega^{\text{post}} = (\omega_\pi^{\text{post}}), \alpha^1+t^1, \alpha^2+t^2\right] \text{ and } \omega_\pi^{\text{post}} \propto p_\pi(t^1|t_\pi^1,y)\, p_\pi(t^2|t_\pi^2,y)\, p_\pi(t_\pi^1,t_\pi^2|y)\, \omega$$

Notice in DDM, we restricted $\beta = \alpha^1+\alpha^2$.

□

**3. Numerical Experiments.**

3.1. *Synthetic Data.* In this section we look more closely at the synthetic data generated using `splatter`. We use PCA plots to show the subtle changes underlying each subtype of simulated data and we demonstrate consistency of estimated distributional changes based on `scDDboost` and Wasserstein distance. Finally, ROC curves illustrate that `scDDboost` has favorable operating characteristics.

We first look at the PCA plots of the simulated data (Supplementary Figure S3, S4, S5). For $K = 7$ and 12, in each scenario there were some subtypes nested in the 2d PCA projection and the distributional change of transcripts becomes difficult to detect. `scDDboost` benefits from the compositional structure and is more sensitive to those subtle changes.

We observed consistent measurements of distributional change by `scDDboost` and Wasserstein distance (Supplementary Figure S6). Lower probabilities of equivalent distributed are associated with bigger distances.

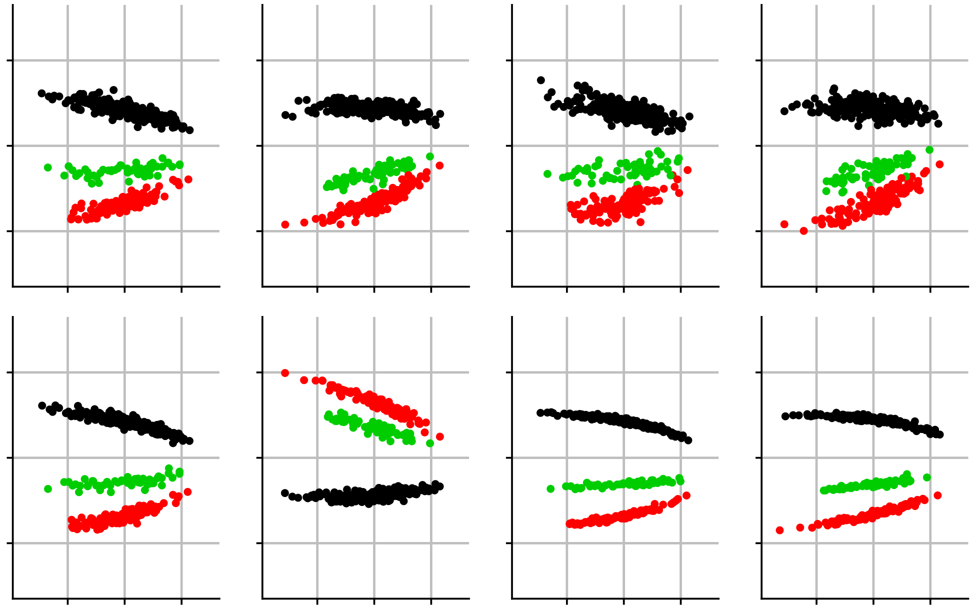

Supplementary Figure S3: First two principal components of transcripts under different parameters for simulated data. Horizontal axis refers to first component, vertical axis refers to second component. Different parameters resulted in different degree of separation of subtypes. We have 4 different settings for hyper-parameters of simulation, each setting has 2 replicates. From left to right, the associated hyper-parameters are (0.1,0.4), (-0.1,0.3), (0.3,0.5), (-0.1,1). Here we have 3 subtypes

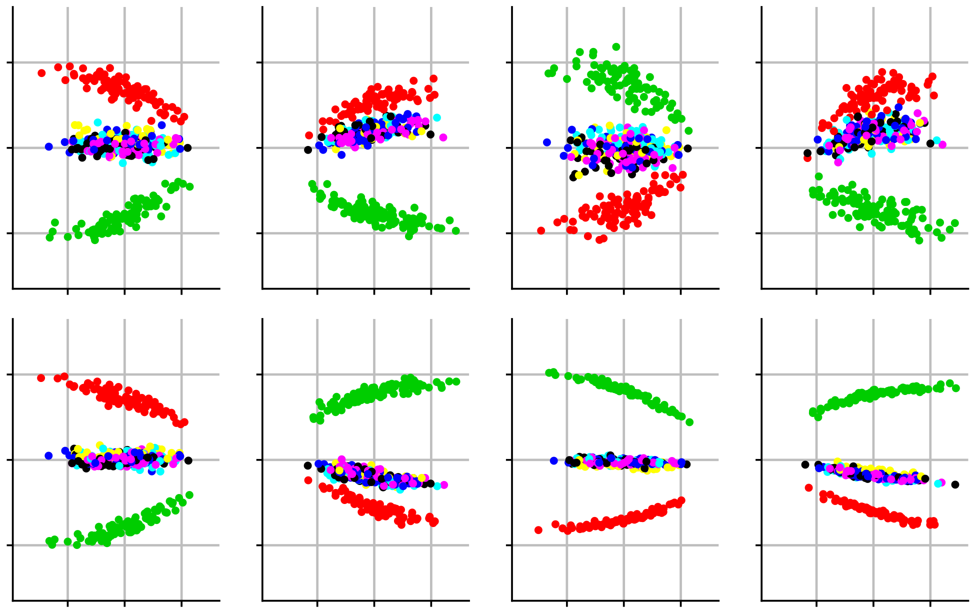

Supplementary Figure S4: Similar plots as Supplementary Figure S3, but for 7 subtypes

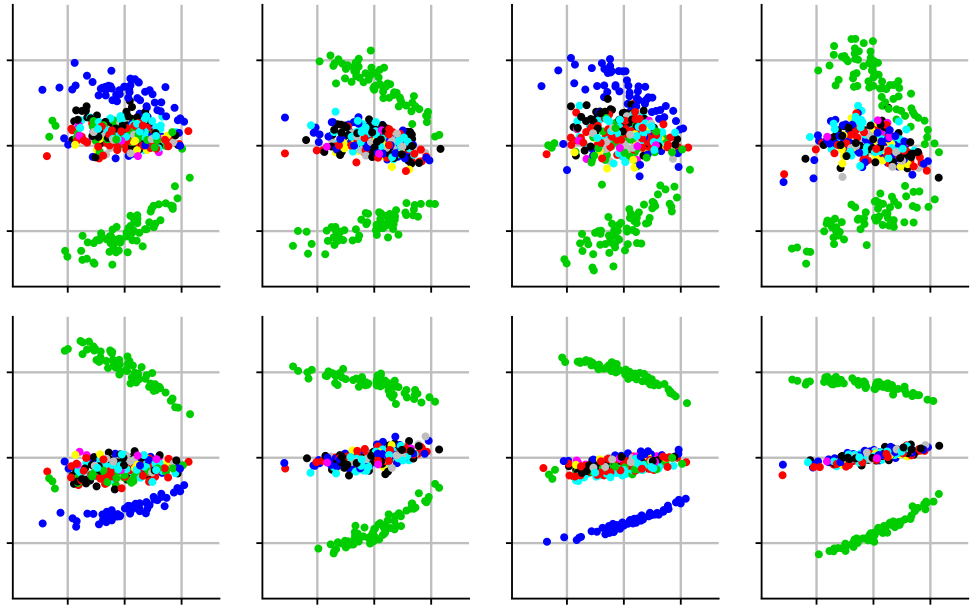

**Supplementary Figure S5:** Similar plots as Sumpplementary Figure S3, but for 12 subtypes

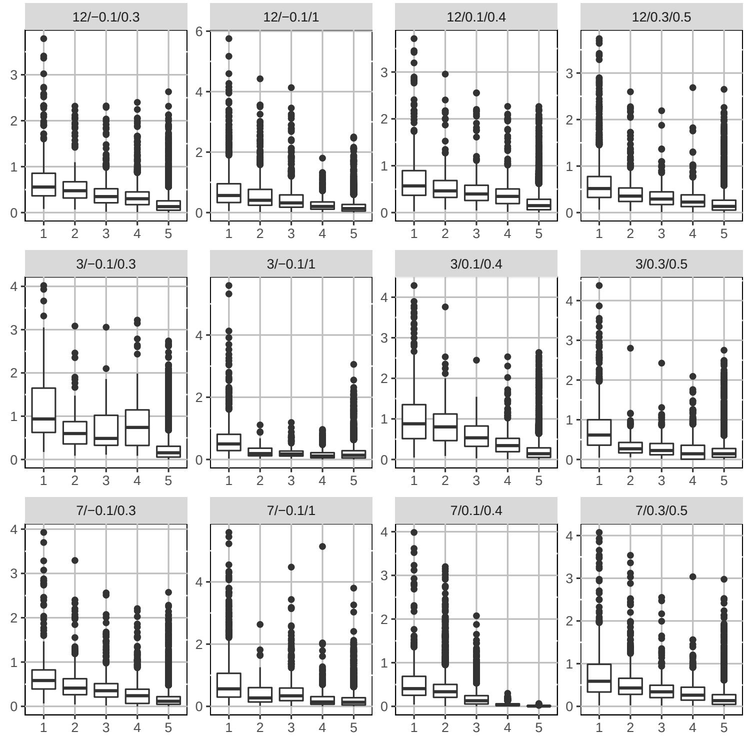


**Supplementary Figure S6:** $P(ED_g|X,y)$ given by scDDboost (horizontal) versus empirical Wasserstein distance (vertical). Genes associated with boxes from left to right having $P(ED_g|X,y)$ range from 0 - 0.2, 0.2 - 0.4, 0.4 - 0.6, 0.6 - 0.8, 0.8 - 1. For simulation cases with parameters in the format: number of clusters / shape / scale

ROC curves for the simulated data in Supplementary Figure S7. Each sub-figure is averaged over two replicates under the same parameters setting. scDDboost tends to outperform other methods .

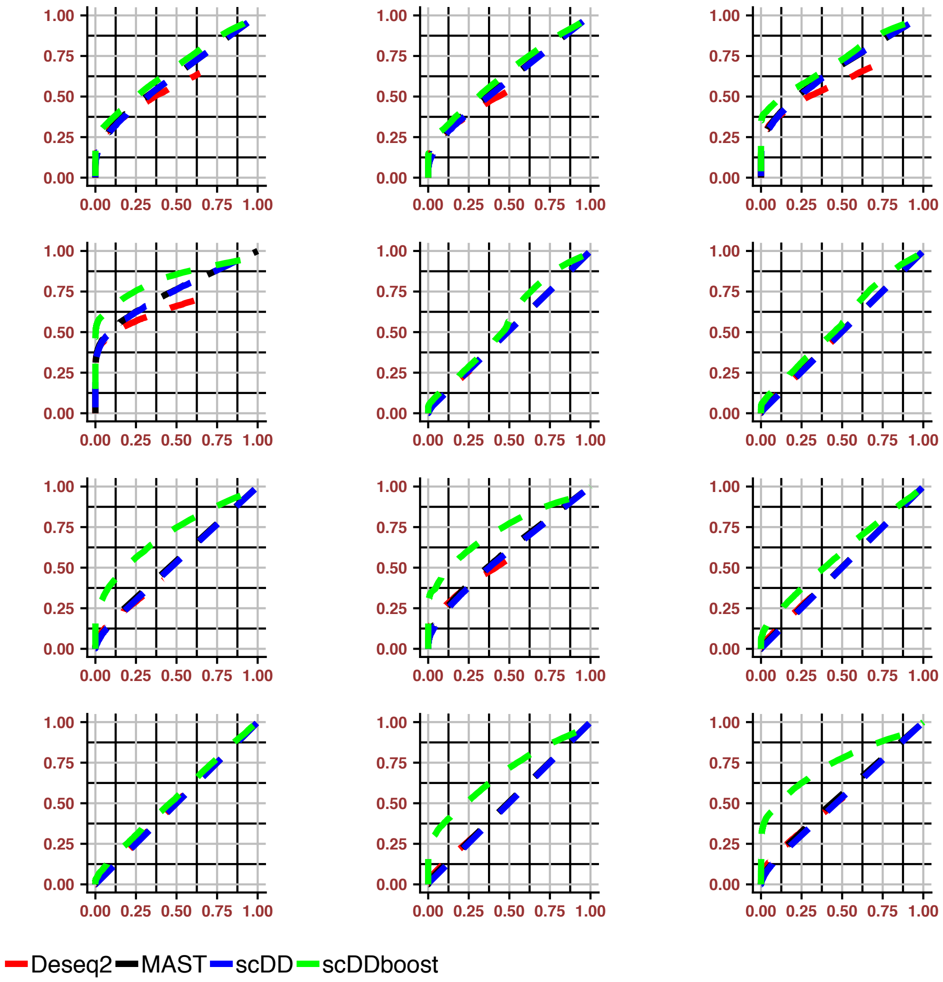


**Supplementary Figure S7:** Roc curve of the 12 simulation settings, under each setting, TPR and FPR are averaged over two replicates, generally `scDDboost` performs better than other methods

3.2. *Empirical Study.* In this section, we provide details of the empirical datasets and also demonstrate consistency to Wasserstein distance on one dataset FUCCI (Leng et al., 2015).

**Data sets** Details for the datasets used in the empirical studies with the estimated number of subtypes $K$ are shown in Supplementary Table S1.

| Data set | Conditions | Number of cells/condition | Organism | Ref | $K$ |
|---|---|---|---|---|---|
| GSE94383 | 0 min unstim vs 75min stim | 186,145 | human | (Lane et al., 2017) | 9 |
| GSE48968-GPL13112 | BMDC (2h LPS stimulation) vs 6h LPS | 96,96 | mouse | (Shalek et al., 2014) | 4 |
| GSE52529 | T0 vs T72 | 69,74 | human | (Trapnell et al., 2014) | 7 |
| GSE74596 | NKT1 vs NTK2 | 46,68 | mouse | (Engel et al., 2016) | 7 |
| EMTAB2805 | G1 vs G2M | 95,96 | mouse | (Buettner et al., 2015) | 6 |
| GSE71585-GPL13112 | Gad2tdTpositive vs Cux2tdTnegative | 80,140 | mouse | (Tasic et al., 2016) | 4 |
| GSE64016 | G1 vs G2 | 91,76 | human | (Leng et al., 2015) | 6 |
| GSE79102 | patient1 vs patient2 | 51, 89 | human | Kiselev et al. (2017) | 4 |
| GSE45719 | 16-cell stage blastomere vs mid blastocyst cell | 50, 60 | mouse | (Deng et al., 2014) | 4 |
| GSE63818 | Primordial Germ Cells, develop- mental stage: 7 week gestation vs Somatic Cells, developmental stage: 7 week gestation | 40,26 | mouse | (Guo et al., 2015) | 6 |
| GSE75748 | DEC vs EC | 64, 64 | human | (Chu et al., 2016) | 5 |
| GSE84465 | neoplastic cells vs non-neoplastic cells | 1000, 1000 | human | (Darmanis et al., 2017) | 9 |

Supplementary Table S1
*Datasets used for empirical study*

For the first 11 datasets in Supplementary Table S1 we use all the cells within that condition under same batch. The last one is the largest dataset we explored containing 3589 cells and comparing neoplastic cells (1091 cells) vs non-neoplastic cells (2498 cells). We randomly sampled 1000 cells from each condition, because it takes a lot of time for `DESeq` and `scDD` to compute when using all the samples and we conjecture that 1000 cells each condition would be enough to represent the heterogeneity. We run the comparison on those subsamples instead and found `DESeq` identified significantly smaller numbers of positives than other methods. It is intuitive that we are more likely to encounter subtle changes when we have large samples, and only considering mean shifts would have limited power.

We also observed consistent distributional change measurements by `scDDboost` and Wasserstein distance (Supplementary Figure S8).

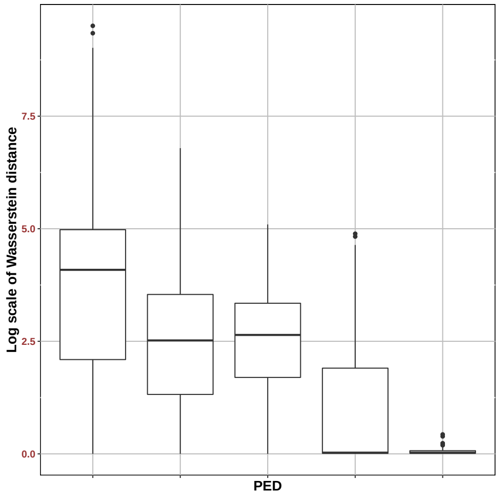


**Supplementary Figure S8:** $P(ED_g|X,y)$ given by scDDboost versus empirical Wasserstein distance. Genes associated with boxes from left to right having $P(ED_g|X,y)$ range from 0 - 0.2, 0.2 - 0.4, 0.4 - 0.6, 0.6 - 0.8, 0.8 - 1, data used: FUCCI

.

Datasets used for generating the Null cases are shown in Supplementary Table S2.

| Data set | Conditions | Number of cells/condition | Organism |
|---|---|---|---|
| GSE63818null | 7 week gestation | 20,20 | mouse |
| GSE75748null | DEC | 32, 32 | human |
| GSE94383null | T0 | 93, 93 | human |
| GSE48968-GPL13112null | BMDC (2h LPS stimulation) | 48,48 | mouse |
| GSE74596null | NKT1 | 23,23 | mouse |
| EMTAB2805null | G1 | 48,48 | mouse |
| GSE71585-GPL13112null | Gad2tdTpositive | 40,40 | mouse |
| GSE64016null | G1 | 46,45 | human |
| GSE79102null | patient1 | 26, 25 | human |

Supplementary Table S2

*Datasets used for null cases, as cells are coming from same biological condition, there should not be any differential distributed genes, any positive call is false positive*

3.3. *Robustness.* In this section, we demonstrate change of the posterior probability of DD under different $K$, and also the robustness provided by random weighting. We also give an example where very large $K$ inflates FDR.

The number of subtypes $K$ is an important parameter. Taking $K$ too small may end up underfitting such that cells within same subtype can still be very different, the mean expression change among subtypes is incapable to capture the marginal distribution change. This would lead to reduced power. Too large $K$ may end up overfitting such that two subtypes can be very similar. Given that we have a fixed number of cells, allowing more clusters will not only increase the burden of computation but decrease the certainty of our inference on DE pattern. Empirically we find that taking $K \leq 10$ is often sufficient (Supplementary Table S1). In any case, we note here that $K$ affects the posterior probability of DD (PDD).

To demonstrate the change of PDD over different $K$, we present an example using dataset GSE75748. When we increase $K$, the variance of the differential term $\text{PDD}_{K+1} - \text{PDD}_K$ keeps decreasing and PDD keeps increasing. Our selection criterion (K = 5) happens to choose $K$ such that change between $\text{PDD}_{K+1}$ and $\text{PDD}_K$ is small while not inflating PDD. We generally obtain stable validity score and PDD simultaneously (Supplementary Figure S9). In addition, the random weighting scheme helps by smoothing PDD (Supplementary Figure S10).

`scDDboost` as the potential to lose FDR control if $K$ is not maintained at a sufficiently small value. Supplementary Figure S11 shows what happens as $K$ increases in one case, other factors staying constant. In our simulation study, we note that the validity score method was always conservative, and did not lead to overestimating $K$.

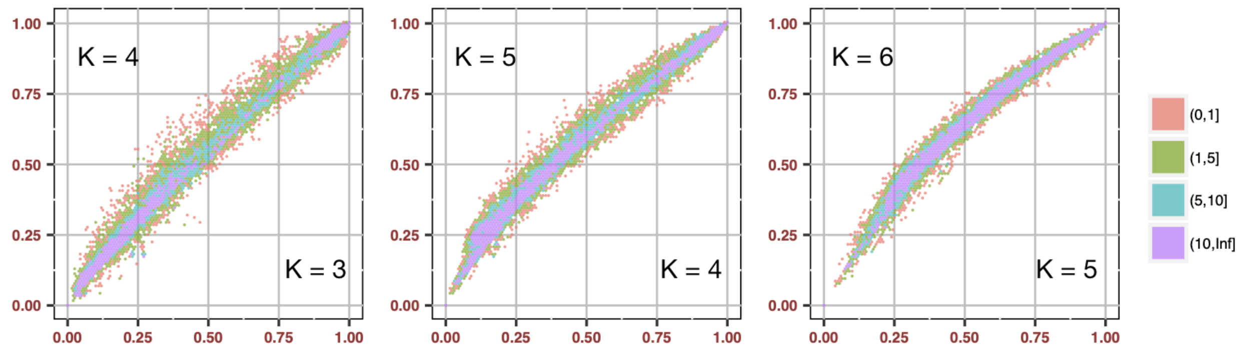


**Supplementary Figure S9:** PDD change under different number of subtypes $K$ for dataset DEC-EC (GSE75748). We select $K = 4$, which also stabilize the PDD.

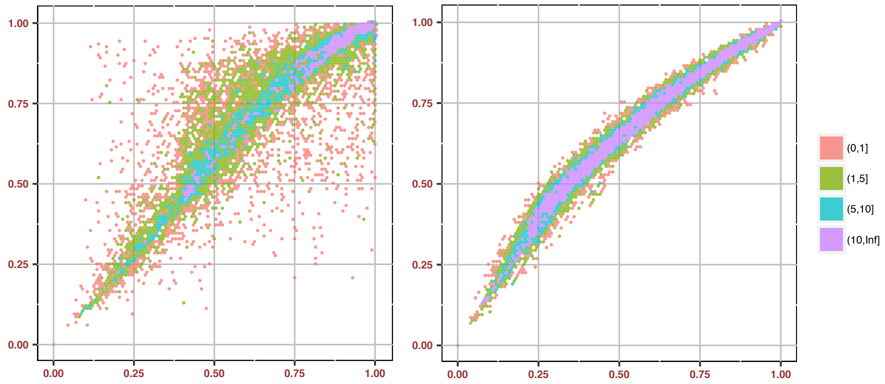


**Supplementary Figure S10:** PDD under $K = 5$ vs. $K = 6$ for dataset DEC-EC (GSE75748). PDD without randomization (left) vs. PDD with randomization (right). `scDDboost` gained robustness through random weighting.

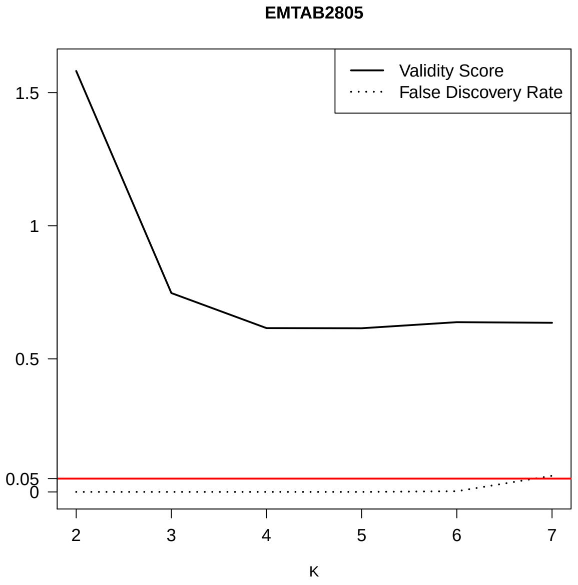


**Supplementary Figure S11:** Under NULL case, using dataset EMTAB2805, when using too big $K$ we may lose FDR control (black dashed line shows proportion of false positive identified by scDDboost under 0.05 threshold, while validity score stabilized after $K > 2$

**4. Posterior consistency.** In this section, we prove Theorem 4 and we discuss a case when condition (12) fails. The density of DDM is computed by product or ratio over several gamma functions. We use a crucial lemma which gives us an approximation to the gamma function, namely

LEMMA 2. *For* $x \geq 1$, $\frac{x^{x-c}}{e^{x-1}} \leq \Gamma(x) \leq \frac{x^{x-1/2}}{e^{x-1}}$, *where* $c = 0.577215...$ *is the Euler-Mascheroni constant.*

PROOF OF LEMMA 2. By (Li and ping Chen, 2007), we have $\frac{x^{x-c}}{e^{x-1}} \leq \Gamma(x) \leq \frac{x^{x-1/2}}{e^{x-1}}$ for $x > 1$ and now we added the case when $x = 1, \Gamma(x) = 1$ so that both sides will include the equality case. □

We have another lemma.

LEMMA 3. *If* $(\phi, \psi) \in A_{\pi_1} \cap A_{\pi_2}$, *follow the conditions in Theorem 1 then*

$$\frac{\omega_{\pi_1}^{post}}{\omega_{\pi_2}^{post}} \xrightarrow[n\to\infty]{a.s.} 0 \quad \text{if } N(\pi_1) < N(\pi_2)$$

PROOF OF LEMMA 3. Recall $\omega_\pi^{\text{post}} \propto p_\pi(t^1|t_\pi^1, y)\, p_\pi(t^2|t_\pi^2, y)\, p_\pi(t_\pi^1, t_\pi^2|y)\, \omega_\pi$ and RHS $= g(\pi, \alpha, \beta, n_1, n_2) f(\pi, t^1, t^2, \alpha, \beta)$ and $\frac{\omega_{\pi_1}^{\text{post}}}{\omega_{\pi_2}^{\text{post}}} = \frac{g(\pi_1,\alpha,\beta,n_1,n_2)}{g(\pi_2,\alpha,\beta,n_1,n_2)} \frac{f(\pi_1,t^1,t^2,\alpha,\beta)}{f(\pi_2,t^1,t^2,\alpha,\beta)}$ where

$$g(\pi, t^1, t^2, \alpha, \beta) = [\prod_{j=1}^{2}\prod_{b\in\pi} \frac{\Gamma(\Sigma_{k\in b}\alpha_k^j)}{\prod_{k\in b}\Gamma(\alpha_k^j)}] \frac{\Gamma(n_1+1)\Gamma(n_2+1)}{\prod_{b\in\pi}\Gamma(\beta_b)} \frac{\Gamma(\Sigma_{b\in\pi}\beta_b)}{\Gamma(n_1+n_2+\Sigma_{b\in\pi}\beta_b)}$$

$$f(\pi, t^1, t^2, \alpha, \beta) = [\prod_{j=1}^{2}\prod_{b\in\pi} \frac{1}{\prod_{k\in b}\Gamma(t_k^j+1)} \frac{\prod_{k\in b}\Gamma(\alpha_k^j + t_k^j)}{\Gamma(t_b^j + \Sigma_{k\in b}\alpha_k^j)}] \prod_{b\in\pi}\Gamma(\beta_b + t_b^1 + t_b^2)$$

For notational simplicity, we use the abbreviation $g(\pi), f(\pi)$ to substitute $g(\pi, \alpha, \beta, n_1, n_2), f(\pi, t^1, t^2, \alpha, \beta)$. We take log on $\frac{\omega_{\pi_1}^{\text{post}}}{\omega_{\pi_2}^{\text{post}}}$, denote it as LR. $LR = \ln g(\pi_1) - \ln g(\pi_2) + \ln f(\pi_1) - \ln f(\pi_2)$. Denote $C(\pi_1, \pi_2, \alpha, \beta) = \ln g(\pi_1) - \ln g(\pi_2)$, $C(\pi_1, \pi_2, \alpha, \beta)$ does not change with sample size $n_1, n_2$ and is a constant determined by partition $\pi_1, \pi_2$ and hyper parameters $\alpha, \beta$. For further convenience of notation let $h(x) = \ln\Gamma(x)$ and $\gamma_b^j = \Sigma_{k\in b}\alpha_k^j$. Denote $R(\pi_1, \pi_2, t^1, t^2, \alpha, \beta) = \ln f(\pi_1) - \ln f(\pi_2)$. And removing the common part of $f(\pi_1)$ and $f(\pi_2)$, we have

$$R(\pi_1, \pi_2, t^1, t^2, \alpha, \beta) = d(\pi_1, t^1, t^2, \alpha, \beta) - d(\pi_2, t^1, t^2, \alpha, \beta)$$

where

$$d(\pi, t^1, t^2, \alpha, \beta) = \sum_{b\in\pi} h(\beta_b + t_b^1 + t_b^2) - \sum_{j=1}^{2}\sum_{b\in\pi} h(t_b + \gamma_b^j)$$

Recall $\beta_b = \gamma_b^1 + \gamma_b^2$ and from Lemma 2, $(x-c)\ln(x) - x + 1 \le h(x) \le (x-1/2)\ln(x) - x + 1$ we have

(3)

$$d(\pi, t^1, t^2, \alpha, \beta) \ge \sum_{b\in\pi} (\beta_b + t_b^1 + t_b^2 - c)\ln(\beta_b + t_b^1 + t_b^2) - \sum_{j=1}^{2}\sum_{b\in\pi} (t_b^j + \gamma_b^j - 1/2)\ln(t_b^j + \gamma_b^j) + N(\pi)$$

(4)

$$d(\pi, t^1, t^2, \alpha, \beta) \le \sum_{b\in\pi} (\beta_b + t_b^1 + t_b^2 - 1/2)\ln(\beta_b + t_b^1 + t_b^2) - \sum_{j=1}^{2}\sum_{b\in\pi} (t_b^j + \gamma_b^j - c)\ln(t_b^j + \gamma_b^j) + N(\pi)$$

$$\text{RHS of (4)} = \Sigma_b\big[(t_b^1 + \gamma_b^1)\ln(1 + \frac{t_b^2 + \gamma_b^2}{t_b^1 + \gamma_b^1}) + (t_b^2 + \gamma_b^2)\ln(1 + \frac{t_b^1 + \gamma_b^1}{t_b^2 + \gamma_b^2})$$
$$+(1-c)\ln(\beta_b + t_b^1 + t_b^2) - 1/2(\ln(1 + \frac{t_b^2 + \gamma_b^2}{t_b^1 + \gamma_b^1}) + \ln(1 + \frac{t_b^1 + \gamma_b^1}{t_b^2 + \gamma_b^2}))\big] + N(\pi)$$

By Taylor expansion at $x = 1$, $\ln(x+1) = \ln 2 + 1/2(x-1) - 1/8(x-1)^2 + g(\xi)(x-1)^3$, where $g(\xi)$ is the reminder term of form $\frac{1}{3(1+\xi)^3}$ for $0 < \xi < x$ For a fixed $n_1, n_2$, we have

$$\begin{aligned}\text{RHS of (4)} = {} & (n_1 + n_2)\ln 2 - \Sigma_{b\in\pi}(1/8(X_b^1 + X_b^2) \\ & + g(\xi_b)(Y_b^1 + Y_b^2)) + T(\pi) + N(\pi)\end{aligned}$$

where $X_b^1 = \frac{(t_b^1 - t_b^2 + \gamma_b^1 - \gamma_b^2)^2}{t_b^1 + \gamma_b^1}$, $X_b^2 = \frac{(t_b^1 - t_b^2 + \gamma_b^1 - \gamma_b^2)^2}{t_b^2 + \gamma_b^2}$, $Y_b^1 = \frac{(t_b^1 - t_b^2 + \gamma_b^1 - \gamma_b^2)^3}{(t_b^1 + \gamma_b^1)^2}$ , $Y_b^2 = \frac{(t_b^1 - t_b^2 + \gamma_b^1 - \gamma_b^2)^3}{(t_b^2 + \gamma_b^2)^2}$ and $T(\pi) = \Sigma_{b\in\pi}\big[(1-c)\ln(\beta_b + t_b^1 + t_b^2) - 1/2(\ln(1 + \frac{t_b^2 + \gamma_b^2}{t_b^1 + \gamma_b^1}) + \ln(1 + \frac{t_b^1 + \gamma_b^1}{t_b^2 + \gamma_b^2}))\big]$
Similarly

$$\begin{aligned}\text{RHS of (5)} = {} & (n_1 + n_2)\ln 2 - \Sigma_{b\in\pi}(1/8(X_b^1 + X_b^2) \\ & + g(\xi_b)(Y_b^1 + Y_b^2)) + U(\pi) + N(\pi)\end{aligned}$$

$U(\pi) = \Sigma_{b\in\pi}\big[(2c - 1/2)\ln(\beta_b + t_b^1 + t_b^2) - c(\ln(1 + \frac{t_b^2 + \gamma_b^2}{t_b^1 + \gamma_b^1}) + \ln(1 + \frac{t_b^1 + \gamma_b^1}{t_b^2 + \gamma_b^2}))\big]$

Using above inequalities, we have

$$\begin{aligned}R(\pi_1, \pi_2, t^1, t^2, \alpha, \beta) \le {} & U(\pi_1) - T(\pi_2) - 1/8(\Sigma_{b\in\pi_1}(X_b^1 + X_b^2) - \Sigma_{b\in\pi_2}(X_b^1 + X_b^2)) \\ & + \Sigma_{b\in\pi_1} g(\xi_b)(Y_b^1 + Y_b^2) - \Sigma_{b\in\pi_2} g(\xi_b)(Y_b^1 + Y_b^2)\end{aligned}$$

$Y_b^j = \frac{((t_b^1 - t_b^2 + \gamma_b^1 - \gamma_b^2)/\sqrt{n}))^3/\sqrt{n}}{((t_b^j + \gamma_b^j)/n)^2}$, by LLN the denominator goes to a constant and by CLT in the numerator $(t_b^1 - t_b^2 + \gamma_b^1 - \gamma_b^2)/\sqrt{n} \to (t_b^1 - t_b^2)/\sqrt{n} \to \sqrt{n}[(t_b^1/n - \Phi_b) - (t_b^2/n -$

$\Psi_b)]$, which converges to a normally distributed random variable when $\Phi_b = \Psi_b$. So $Y_b^j$ is $o_p(1)$. Similarly, $X_b^j = \frac{((t_b^1 - t_b^2 + \gamma_b^1 - \gamma_b^2)/\sqrt{n})^2}{t_b^j + \gamma_b^j / n}$ is asymptotically gamma ($\chi$-square) distributed. $g(\xi_b)$ has bounded variance, $U(\pi_1) - T(\pi_2) = -\ln(n)$ if $N(\pi_2) < N(\pi_1)$ as $\ln(\beta_b + t_b^1 + t_b^2) - \ln(\beta_{b'} + t_{b'}^1 + t_{b'}^2) = \ln(\frac{\beta_b + t_b^1 + t_b^2}{n}) - \ln(\frac{\beta_{b'} + t_{b'}^1 + t_{b'}^2}{n}) \to O(1) \quad a.s.$, which completes the proof.

□

Proof of Theorem 4. Recall $\sum_{\pi \in \Pi} \omega_\pi^{\text{post}} = 1$ and $P(A_\pi | y, z) = \sum_{\tilde{\pi} \in \Pi} \omega_{\tilde{\pi}}^{\text{post}} 1[\tilde{\pi} \text{ refines } \pi]$. If $(\phi, \psi) \notin$ Q, for all the $A_\pi$ covers $(\phi, \psi)$ there is one finest $\pi^*$ with the largest $N(\pi^*)$ and every other $\pi$ that $(\phi, \psi) \in A_\pi$ is coarser than $\pi^*$. Theorem 4 now follows by Lemma 3.

□

Under some choices of $(\phi, \psi)$, condition (12) could fail.

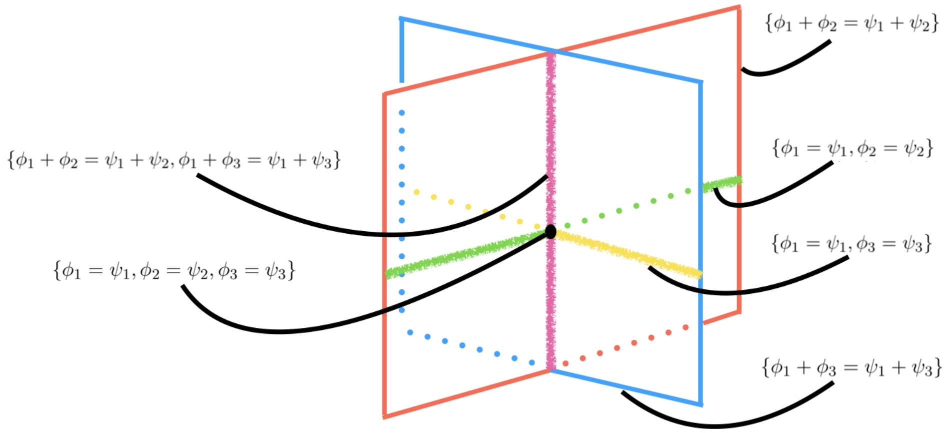


**Supplementary Figure S12:** Four subtypes of cells, simplexes of $(\phi, \psi)$ satisfying different constraints.

In Supplementary Figure S12, there are four subtypes, the rectangle with magenta boundary is a simplex $A_{\pi_1} = \{(\phi, \psi) : \phi_1 + \phi_2 = \psi_1 + \psi_2\}$, the rectangle with blue boundary is another simplex $A_{\pi_2} = \{(\phi, \psi) : \phi_1 + \phi_3 = \psi_1 + \psi_3\}$. The green line refers to $A_{\pi_3} = \{(\phi, \psi) : \phi_1 = \psi_1, \phi_2 = \psi_2\}$, the yellow line refers to $A_{\pi_4} = \{(\phi, \psi) : \phi_1 = \psi_1, \phi_3 = \psi_3\}$. the purple line refers to $O = \{(\phi, \psi) : \phi_1 + \phi_2 = \psi_1 + \psi_2, \phi_1 + \phi_3 = \psi_1 + \psi_3\}$, which is the intersection of $A_{\pi_1}$ and $A_{\pi_2}$, and finally the black dot which is the intersection of those three lines refers to the simplex with finest partitions, $\phi_i = \psi_i, \forall i = 1, \cdots, 4$. When $(\phi, \psi)$ is from the purple line except the black dot, condition (12) would fail as there is not a finest $\pi^*$ of $H(\phi, \psi)$. This may be of theoretical interest, but the practical implications of this finding are negligible as further computations have demonstrated.

### References.


Buettner, F., Natarajan, K. N., Casale, F. P., Proserpio, V., Scialdone, A., Theis, F. J., Teichmann, S. A., Marioni, J. C. and Stegle, O. (2015). Computational analysis of cell-to-cell heterogeneity in single-cell RNA-sequencing data reveals hidden subpopulations of cells. *Nature Biotechnology* **33** 155 EP -.

Chu, L.-F., Leng, N., Zhang, J., Hou, Z., Mamott, D., Vereide, D. T., Choi, J., Kendziorski, C., Stewart, R. and Thomson, J. A. (2016). Single-cell RNA-seq reveals novel regulators of human embryonic stem cell differentiation to definitive endoderm. *Genome Biology* **17** 173. .

Dahl, D. B. (2009). Modal clustering in a class of product partition models. *Bayesian Anal.* **4** 243–264.

Darmanis, S., Sloan, S. A., Croote, D., Mignardi, M., Chernikova, S., Samghababi, P., Zhang, Y., Neff, N., Kowarsky, M., Caneda, C., Li, G., Chang, S. D., Connolly, I. D., Li, Y., Barres, B. A., Gephart, M. H. and Quake, S. R. (2017). Single-Cell RNA-Seq Analysis of Infiltrating Neoplastic Cells at the Migrating Front of Human Glioblastoma. *Cell reports* **21** 1399–1410.

Deng, Q., Ramsköld, D., Reinius, B. and Sandberg, R. (2014). Single-Cell RNA-Seq Reveals Dynamic, Random Monoallelic Gene Expression in Mammalian Cells. *Science* **343** 193–196.

Engel, I., Seumois, G., Chavez, L., Samaniego-Castruita, D., White, B., Chawla, A., Mock, D., Vijayanand, P. and Kronenberg, M. (2016). Innate-like functions of natural killer T cell subsets result from highly divergent gene programs. *Nature Immunology* **17** 728 EP -.

Guo, F., Yan, L., Guo, H., Li, L., Hu, B., Zhao, Y., Yong, J., Hu, Y., Wang, X., Wei, Y., Wang, W., Li, R., Yan, J., Zhi, X., Zhang, Y., Jin, H., Zhang, W., Hou, Y., Zhu, P., Li, J., Zhang, L., Liu, S., Ren, Y., Zhu, X., Wen, L., Gao, Y. Q., Tang, F. and Qiao, J. (2015). The Transcriptome and DNA Methylome Landscapes of Human Primordial Germ Cells. *Cell* **161** 1437–1452.

Jara, A., Hanson, T., Quintana, F., Mller, P. and Rosner, G. (2011). DPpackage: Bayesian Semi- and Nonparametric Modeling in R. *Journal of Statistical Software, Articles* **40** 1–30.

Kiselev, V. Y., Kirschner, K., Schaub, M. T., Andrews, T., Yiu, A., Chandra, T., Natarajan, K. N., Reik, W., Barahona, M., Green, A. R. and Hemberg, M. (2017). SC3: consensus clustering of single-cell RNA-seq data. *Nature Methods* **14** 483 EP -.

Lane, K., Van Valen, D., DeFelice, M. M., Macklin, D. N., Kudo, T., Jaimovich, A., Carr, A., Meyer, T., Pe'er, D., Boutet, S. C. and Covert, M. W. (2017). Measuring Signaling and RNA-Seq in the Same Cell Links Gene Expression to Dynamic Patterns of NF-B Activation. *Cell Systems* **4** 458–469.e5.

Leng, N., Dawson, J. A., Thomson, J. A., Ruotti, V., Rissman, A. I., Smits, B. M. G., Haag, J. D., Gould, M. N., Stewart, R. M. and Kendziorski, C. (2013). EBSeq: an empirical Bayes hierarchical model for inference in RNA-seq experiments. *Bioinformatics* **29** 1035-1043.

Leng, N., Chu, L.-F., Barry, C., Li, Y., Choi, J., Li, X., Jiang, P., Stewart, R. M., Thomson, J. A. and Kendziorski, C. (2015). Oscope identifies oscillatory genes in unsynchronized single-cell RNA-seq experiments. *Nature Methods* **12** 947 EP -.

Li, X. and ping Chen, C. (2007). Inequalities for the gamma function. In *2007), Art. 28. [ONLINE: http://jipam.vu.edu.au/article.php?sid=842*.

Ray, S. and Turi, R. H. (2000). Determination of Number of Clusters in K-Means Clustering and Application in Colour Image Segmentation.

Shalek, A. K., Satija, R., Shuga, J., Trombetta, J. J., Gennert, D., Lu, D., Chen, P., Gertner, R. S., Gaublomme, J. T., Yosef, N., Schwartz, S., Fowler, B., Weaver, S., Wang, J., Wang, X., Ding, R., Raychowdhury, R., Friedman, N., Hacohen, N., Park, H., May, A. P. and Regev, A. (2014). Single-cell RNA-seq reveals dynamic paracrine control of cellular variation. *Nature* **510** 363 EP -.

Tasic, B., Menon, V., Nguyen, T. N., Kim, T. K., Jarsky, T., Yao, Z., Levi, B., Gray, L. T., Sorensen, S. A., Dolbeare, T., Bertagnolli, D., Goldy, J., Shapovalova, N., Parry, S., Lee, C., Smith, K., Bernard, A., Madisen, L., Sunkin, S. M., Hawrylycz, M., Koch, C. and Zeng, H. (2016). Adult mouse cortical cell taxonomy revealed by single cell transcriptomics. *Nature Neuroscience* **19** 335 EP -.

Trapnell, C., Cacchiarelli, D., Grimsby, J., Pokharel, P., Li, S., Morse, M., Lennon, N. J., Livak, K. J., Mikkelsen, T. S. and Rinn, J. L. (2014). The dynamics and regulators of cell fate decisions are revealed by pseudotemporal ordering of single cells. *Nature biotechnology* **32** 381–386.
